# *STAT3*-dependent systems-level analysis reveals *PDK4* as an independent predictor of biochemical recurrence in prostate cancer

**DOI:** 10.1101/770701

**Authors:** Monika Oberhuber, Matteo Pecoraro, Mate Rusz, Georg Oberhuber, Maritta Wieselberg, Peter Haslinger, Elisabeth Gurnhofer, Jan Pencik, Robert Wiebringhaus, Michaela Schlederer, Theresa Weiss, Margit Schmeidl, Andrea Haitel, Marc Brehme, Wolfgang Wadsak, Johannes Griss, Thomas Mohr, Alexandra Hofer, Anton Jäger, Gerda Egger, Jürgen Pollheimer, Gunda Koellensperger, Matthias Mann, Brigitte Hantusch, Lukas Kenner

**Affiliations:** Department of Pathology; Medical University of Vienna; Vienna, Vienna, 1090; Austria; CBmed - Center for Biomarker Research in Medicine GmbH; Graz, Styria, 8010; Austria; Department of Proteomics and Signal Transduction; Max Planck Institute of Biochemistry; Martinsried, Bavaria, 82152; Germany; Department of Analytical Chemistry; Faculty of Chemistry; University of Vienna; Vienna, Vienna, 1090; Austria; Institute of Inorganic Chemistry; University of Vienna; Vienna, Vienna, 1090; Austria; Patho im Zentrum; St.Pölten, Lower Austria, 3100; Austria; Department of Obstetrics and Gynaecology, Reproductive Biology Unit; Medical University of Vienna; Vienna, Vienna, 1090; Austria; Department of biomedical imaging and Image-guided therapy; Division of Nuclear Medicine; Medical University of Vienna; Vienna, Vienna, 1090; Austria; Department of Dermatology; Medical University of Vienna; Vienna, Vienna, 1090; Austria; Institute of Cancer Research and Comprehensive Cancer Center; Department of Medicine I; Medical University of Vienna; Vienna, Vienna, 1090; Austria; Science Consult DI Thomas Mohr KG; Guntramsdorf, Lower Austria, 2353; Austria; E166 - Institute of Chemical Engineering; Bioprocess Technology; Vienna University of Technology; Vienna, Vienna, 1060; Austria; Ludwig Boltzmann Institute Applied Diagnostics; Vienna, Vienna, 1090; Austria; Christian Doppler Laboratory for Applied Metabolomics; Vienna, Vienna, 1090; Austria; Unit of Pathology of Laboratory Animals; University of Veterinary Medicine Vienna; Vienna, Vienna, 1200; Austria; Ludwig Boltzmann Institute for Cancer Research; Vienna, Vienna, 1090; Austria

## Abstract

Prostate cancer (PCa) has a broad spectrum of clinical behaviour, hence biomarkers are urgently needed for risk stratification. We previously described the protective effect of STAT3 in a prostate cancer mouse model. By utilizing a gene co-expression network in addition to laser microdissected proteomics from human and murine prostate FFPE samples, we describe STAT3-induced downregulation of the TCA cycle/OXPHOS in PCa on transcriptomic and proteomic level. We identify pyruvate dehydrogenase kinase 4 (PDK4), a key regulator of the TCA cycle, as a promising independent prognostic marker in PCa. *PDK4* predicts disease recurrence independent of diagnostic risk factors such as grading, staging and PSA level. Furthermore, *PDK4* expression is causally linked to type 2 diabetes mellitus, which is known to have a protective effect on PCa. We conclude that this effect is related to *PDK4* expression and that *PDK4* loss could serve as a biomarker for PCa with dismal prognosis.

## Introduction

Prostate Cancer (PCa) is the second most frequent cancer and the fifth leading cause of death from cancer in men worldwide (Bray et al., 2018). The diagnosis of PCa is largely based on the histopathological evaluation of biopsies, which are graded by the Gleason score (GSC) (Gleason and Mellinger, 1974). In 2005, the GSC was modified by the International Society of Urological Pathology (ISUP) (Epstein et al., 2005), resulting in the ISUP grade, which ranges from I to V (National Collaborating Centre for Cancer, 2014). PCa shows a wide variety in clinical behaviour, ranging from harmless, indolent tumors to aggressive metastatic disease (Epstein and Lotan, 2014, Sathianathen et al., 2018). As a consequence, treatment following biopsy of the prostate is individualized and based on four main criteria: the amount of tumor in the biopsy, the histological GSC/ISUP grading, clinical staging and – to a lesser extent – the level of prostate specific antigen (PSA) in the serum (National Collaborating Centre for Cancer, 2014). Nonetheless, there is a significant risk of over- and undertreatment (Sathianathen et al., 2018), and additional biomarkers for risk stratification are urgently needed.

Molecular characterization reveals PCa as a highly heterogeneous disease with diverse genetic, epigenetic and transcriptomic alterations (The Cancer Genome Atlas Research Network, 2015, Taylor et al., 2010). As a consequence, there is a strong need to define molecular subgroups of PCa to identify potential targets for treatment. In a previous attempt, our group studied the role of signal transducer and activator of transcription 3 (STAT3) in PCa, which turned out to exert tumor suppressor activities (Pencik et al., 2015). Physiologically, STAT3 plays a central role in signal transduction in a wide range of transcriptional pathways. In cancer, it has been shown to promote stem-cell like characteristics of tumor cells, tumor cell survival and proliferation. STAT3 enhances metastatic potential and immune evasion (Huynh et al., 2019). Owing to these facts, it is generally considered an oncogene. However, others have shown that STAT3 may also have tumor suppressive capabilities (Huynh et al., 2019, Pencik et al., 2015).

In this study, we have analyzed the system-level effects of STAT3 on the transcriptome and proteome of human PCa samples and a previously established (Pencik et al., 2015) *Stat3* PCa mouse model (Figure 1A). For transcriptomic analyses, we used The Cancer Genome Atlas - Prostate Adenocarcinoma (TCGA PRAD) RNA-Seq dataset (The Cancer Genome Atlas Research Network, 2015) and established a gene co-expression network. We found a negative association of *STAT3* expression to the tricarboxylic acid (TCA) cycle / oxidative phosphorylation (OXPHOS) and ribosomal biogenesis. These results were corroborated by findings in shotgun proteomics of laser-microdissected formalin fixed and paraffin embedded (FFPE) PCa material from both human and murine samples. Furthermore, we showed elevated levels of metabolites of the TCA cycle in *PtenStat3^pc-/-^* mice, using a targeted metabolomics approach.

**Figure 1:**
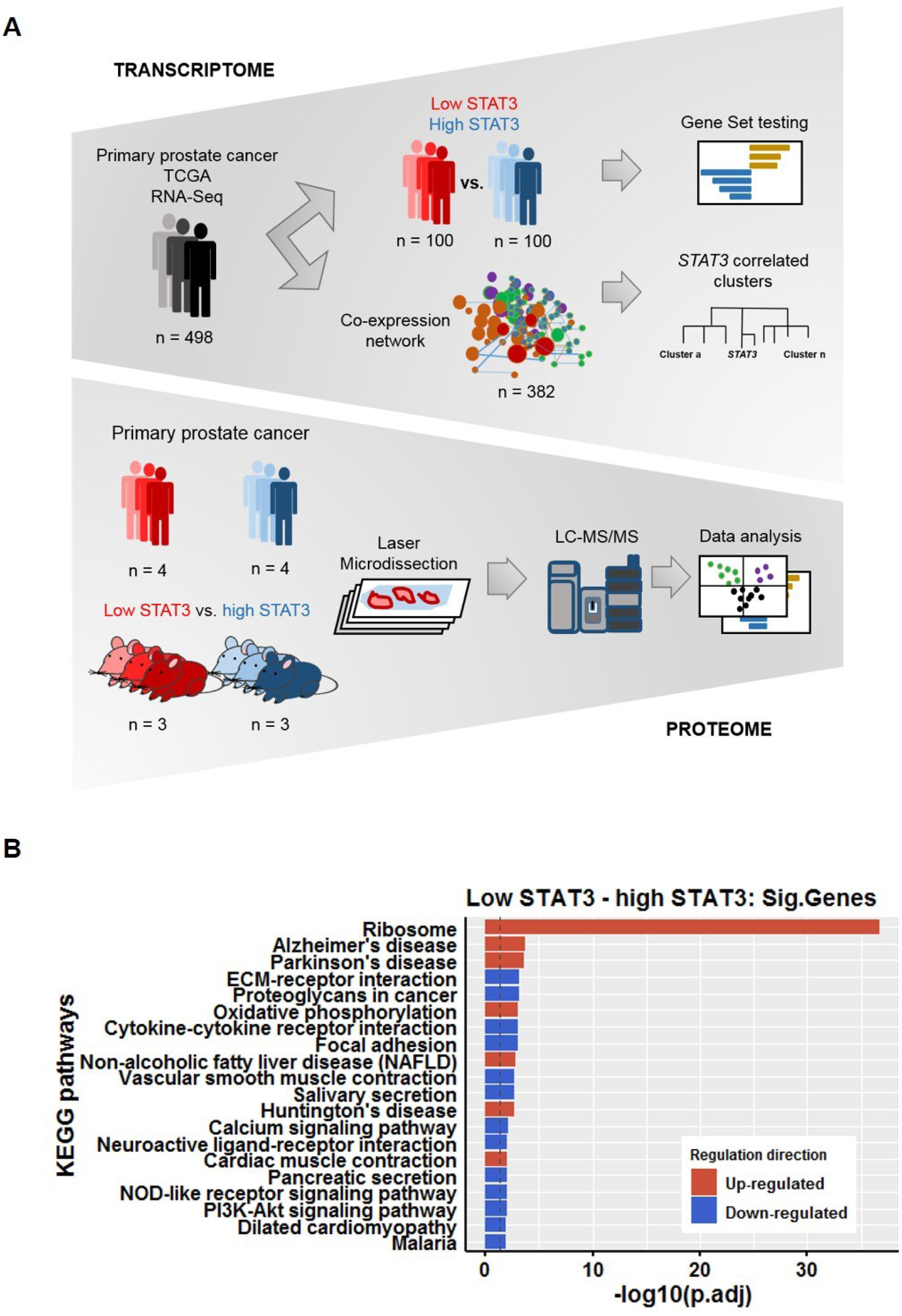
Identification of STAT3 associated pathways in prostate cancer. A. Overview of transcriptomic (top) and proteomic (bottom) analyses. B. Overexpression analysis of enriched KEGG pathways of significantly differentially expressed genes between low STAT3 versus high STAT3 groups in TCGA PRAD. See also Table S1, Figures S1 and S2.

Our experimental results therefore support the notion that STAT3 is an important negative regulator of TCA/OXPHOS (Huynh et al., 2019), thereby influencing aggressiveness of PCa. Furthermore, gene expression of *PDK4*, which inhibits pyruvate oxidation through the TCA cycle and thereby negatively impacts OXPHOS (Zhang et al., 2014), was significantly downregulated in low *STAT3* patients. We show that high *PDK4* expression is significantly associated with a lower risk of biochemical recurrence (BCR), and it predicts disease recurrence independent of ISUP grading in low/intermediate risk primary tumors. In addition, *PDK4* is an independent predictor of BCR compared to ISUP grading and clinical staging, as well as pathological staging and pre-surgical PSA-levels in primary and metastatic tumors, identifying *PDK4* as a promising prognostic marker in PCa.

## Results

### Low *STAT3* expression in primary PCa is associated with increased OXPHOS and ribosomal biosynthesis

In order to gain insight into the effects of *STAT3* expression in primary PCa, we employed two different approaches of analyzing TCGA PRAD RNA-Seq data of 498 patients (Figure 1A).

Firstly, samples were ranked according to *STAT3* expression and split into three groups: “high STAT3” consisted of the 1 - 0.8^th^ quantile (n=100), “low STAT3” of the 0.2^nd^ quantile (n=100) and “medium STAT3” of all samples in between (n=298). We compared low STAT3 to high STAT3 samples and found 1194 genes to be significantly differentially expressed (log-FC ≥ 1, adj. p-value ≤ 0.05, Table S1). Gene set testing using the Ensemble Of Gene Set Enrichment Analyses (EGSEA) method (Alhamdoosh et al., 2017) (Methods) showed gene sets directly associated with STAT3 signaling to be downregulated (Figure S1A-B, Table S1). Interestingly, the hallmark signature gene set for OXPHOS was strongly upregulated, so was the KEGG pathway Ribosome (Table S1). Overexpression analysis of differentially expressed genes showed the ribosome and OXPHOS among upregulated KEGG pathways (Figure 1B, Figure S2A - B), while Gene Ontologies (GO) Cellular Component (CC) showed enrichment of genes coding for ribosomal subunits, RNA metabolism and protein localization, which are involved in translation (Steitz, 2008) (Table S1).

Secondly, we used Weighted Gene Co-Expression Network Analysis (WGCNA) (Methods) (Langfelder and Horvath, 2008, Langfelder and Horvath, 2012) to create a network of co-expressed gene clusters from the entire dataset. Gene clusters consist of groups of genes which are highly interconnected by their high absolute correlation. Gene clusters can be analyzed for common biological motives and for their association with a trait of interest, such as tumor grade, stage or expression of a specific gene. We generated a network from 13,932 genes and 382 patients which resulted in 13 gene clusters that we analyzed for characteristic biological themes by using overexpression analysis (Yu et al., 2012) (Figure 2A and 2B). Some gene clusters indicated specific overexpression of distinct biological motives. Genes in cluster 2 for example, were mainly associated with cellular respiration (mitochondrial respiratory complex assembly, OXPHOS) and RNA splicing. Cluster 3 represented ribosomal translation and protein targeting to the endoplasmic reticulum (ER). Cluster 11 was associated with epigenetic processes (histone and chromatin modification, gene silencing). We subsequently investigated which gene clusters were associated with *STAT3* expression by using the cluster eigengene (= the first principal component, Methods) and found strong correlations for three clusters. While genes in “epigenetic”- cluster 11 (Pearson correlation; ρ = 0.59, adj. p-value = 8e-36) showed a positive correlation with *STAT3* expression, “OXPHOS”- cluster 2 (ρ = -0.67, adj. p-value = 7e-50) and “ribosomal”- cluster 3 (ρ = -0.74, adj. p-value = 1e-65) were negatively correlated (Figure 3A and 3B, Table S2). We also investigated the correlation of gene clusters with clinical traits representing tumor aggressiveness. Clusters correlating with the clinical traits BCR, GSC, and the risk groups pathological tumor (pT) and lymph node (pN) staging showed different correlations than those correlated with *STAT3* expression (Figure 3B).

**Figure 2:**
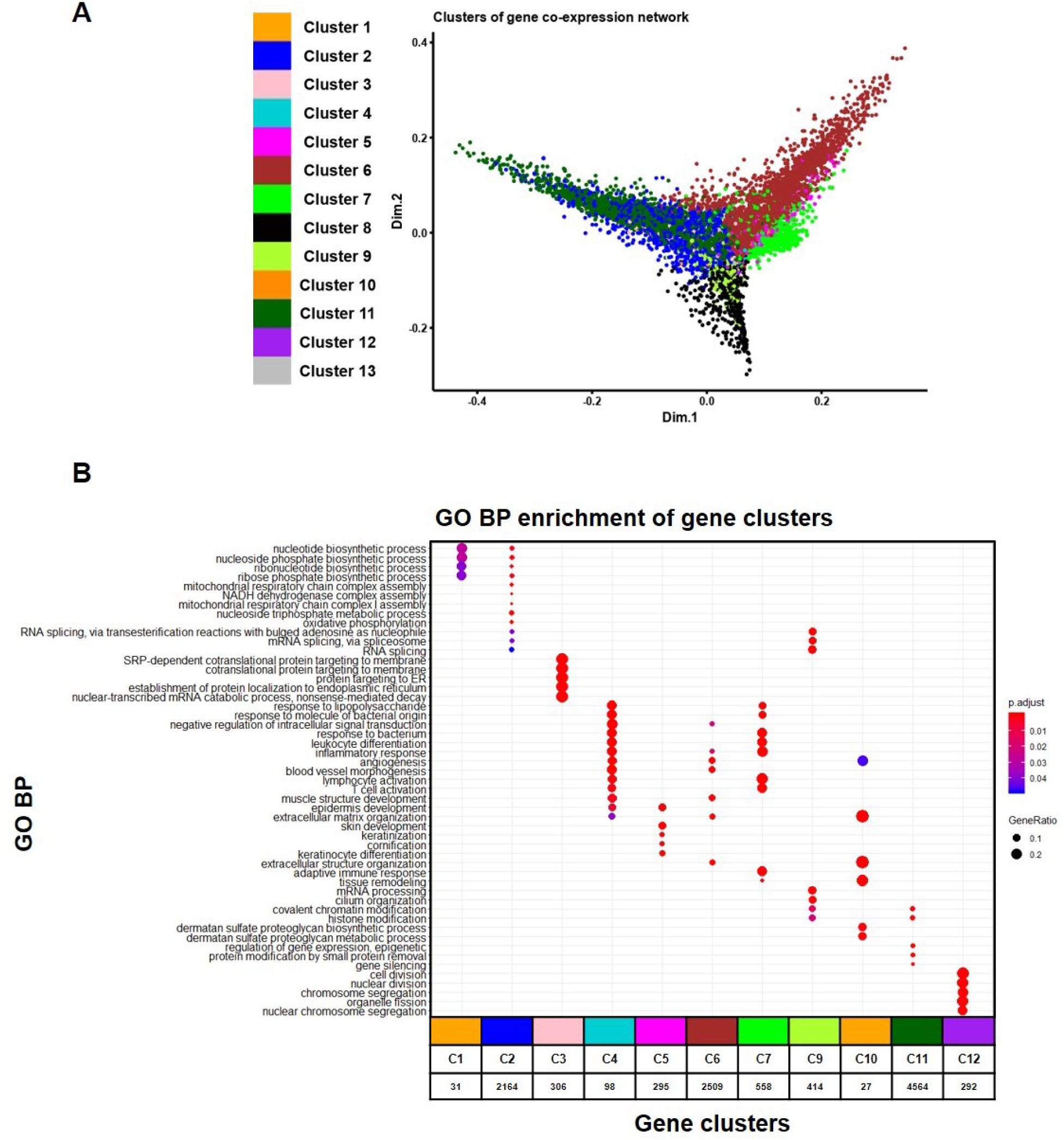
Prostate cancer gene co-expression network shows biological motives of gene clusters. A. Multidimensional scaling (MDS) plot of the prostate cancer gene co-expression network. MDS plot was generated by using the topological overlap matrix (TOM). The topological overlap indicates, whether two genes share co-expression to a similar set of other genes. Colors represent different gene clusters. Genes in a cluster are interconnected by their high absolute correlation. The legend shows gene cluster numbers and respective assigned cluster colors. Smaller clusters may be occluded by larger ones in the MDS plot. B. Biological themes comparison of enriched GO BP terms for all gene clusters. Only clusters shown in the figure contain significantly enriched gene sets. Numbers below clusters indicate the number of genes enriched in GO BPs. Dot color represents significance levels ranging from < 0.01 (= red) to 0.05 (= blue). Dot size represents the gene ratio (number of genes in the cluster significant in the GO term / number of all genes in the cluster). C = cluster, GO = Gene Ontology, BP = Biological Process.

**Figure 3:**
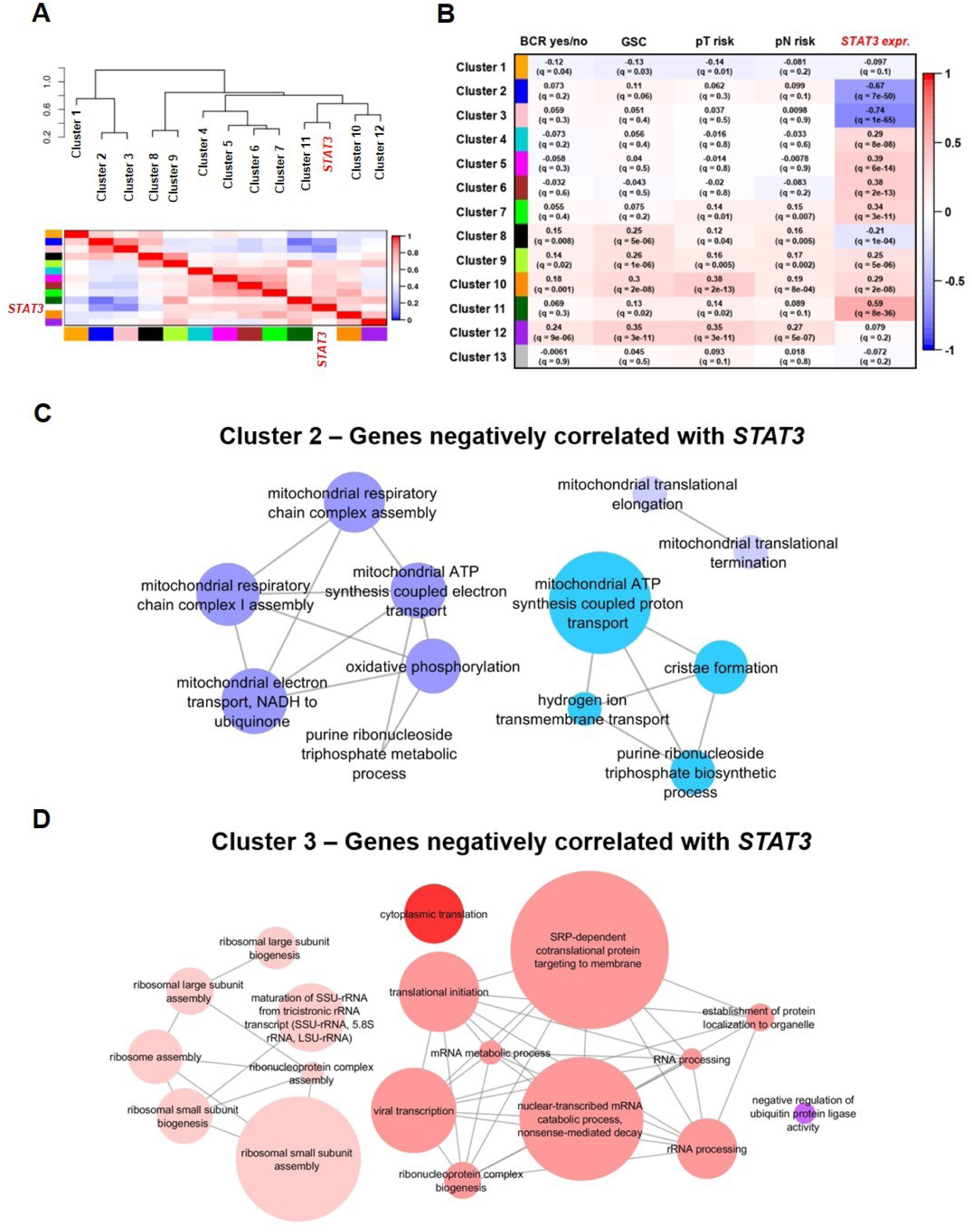
Clusters negatively correlated with *STAT3* are associated with OXPHOS and ribosomal biosynthesis. A. Graphical representation of the network of cluster eigengenes (= their first principal component). Dendrogram and heatmap plots show the relationships between the eigengenes and *STAT3*. Correlations between cluster eigengenes and the trait are indicated by a color bar ranging from red (1) to blue (-1). B. Heatmap showing the correlation of gene cluster eigengenes with traits of interest. Pearson correlation is indicated by colors and values and ranges from 1 (red) to -1 (blue). Adj. p-values (q-values) indicate significance of correlations. BCR = biochemical recurrence, GSC = Gleason Score, pT = pathological tumor staging, pN = pathological lymph node staging, STAT3 expr = *STAT3* gene expression in counts per million (cpm). Low risk = pT2abc, pN0; High risk = pT3-T4, pN1. C. and D. Network representation of enriched GO BP terms of top 50 genes most strongly negatively correlated with *STAT3* (GS ≤ - 0.6, adj. p-value ≤ 0.01) in cluster 2 (C, blue, MM ≥ 0.8, adj. p-value ≤ 0.01) and cluster 3 (D, pink, MM ≥ 0.8, adj. p-value ≤ 0.01), respectively. Node size indicates the percentage of associated genes. Similar colors indicate terms of the same GO group. GS = Gene significance, MM = module membership, GO = Gene Ontology, BP = Biological Process.

To confirm the negative association of *STAT3* with OXPHOS and ribosomal activity, we assessed the correlation of individual genes in cluster 2 and cluster 3 with *STAT3*. For each cluster, we selected the 50 genes that were most strongly negatively correlated with *STAT3* (ρ ≤ - 0.6), while at the same time being highly associated with the respective gene cluster (ρ ≥ 0.8). Those of cluster 2 showed enrichment of GO Biological Process (BP) terms belonging to three groups: Oxidative phosphorylation (45.71%), mitochondrial ATP synthesis coupled protein transport (42.86%) and mitochondrial translational elongation (11.43%) (Figure 3C). In cluster 3, GO BP terms were associated with four groups, consisting of SRP-dependent co-translational protein targeting to membrane (50%), ribosomal small subunit biogenesis (38.89 %), negative regulation of ubiquitin protein ligase activity (5.56%) and cytoplasmic translation (5.56%) (Figure 3D). As a conclusion, both our first and our second analysis suggest a negative correlation of *STAT3* expression to genes associated with both increased OXPHOS and ribosomal activity.

### Proteomics analysis of human FFPE-samples shows high TCA/OXPHOS in low STAT3 PCa

After the investigation of *STAT3*-effects on the gene expression level, we examined its impact on the protein level (Figure 1A). Low and high STAT3 levels were preselected by immunohistochemical (IHC) analyses of STAT3 in patient samples. We conducted shotgun proteomics experiments with FFPE patient material, comparing low STAT3 with high STAT3 PCa and a healthy prostate control group (n= 4 in each group, Methods). To specifically focus on cell autonomous mechanisms, tumor and control material was procured by laser-microdissection (LMD) of prostate epithelial cells, and a label-free quantification (LFQ) approach was used to obtain protein intensities. Samples showed a clear separation of groups after principal component analysis (PCA) (Figure 4A). We identified 2316 proteins on average (1722 - 2610), of which 86 were differentially expressed across all groups (log-FC ≥ 1, adj.p-value < 0.05). Among the 22 proteins we found to be significantly differentially expressed between low STAT3 and high STAT3 groups (log-FC ≥ 1, adj. p-value < 0.05, Figure 4B, Table S3), 13 were mitochondrial proteins. Of interest, succinate dehydrogenase complex iron sulfur subunit B (SDHB) (log-FC = 3.56, adj. p-value = 0.01) and isocitrate dehydrogenase (NADP (+)) 2 (IDH2) (log- FC = 2.31, adj. p-value = 0.0539) were significantly up-regulated. STAT3 could not be detected in this experiment, which may be due to archived FFPE material, which could impede proteomics coverage. Gene set testing between low STAT3 and high STAT3 showed mitochondrial proteins and metabolic processes to be upregulated (Table S3). Consistent with the results of our TCGA analysis, several metabolic KEGG pathways, among them the TCA cycle and OXPHOS (Figure 4C, Table S3), were upregulated.

**Figure 4:**
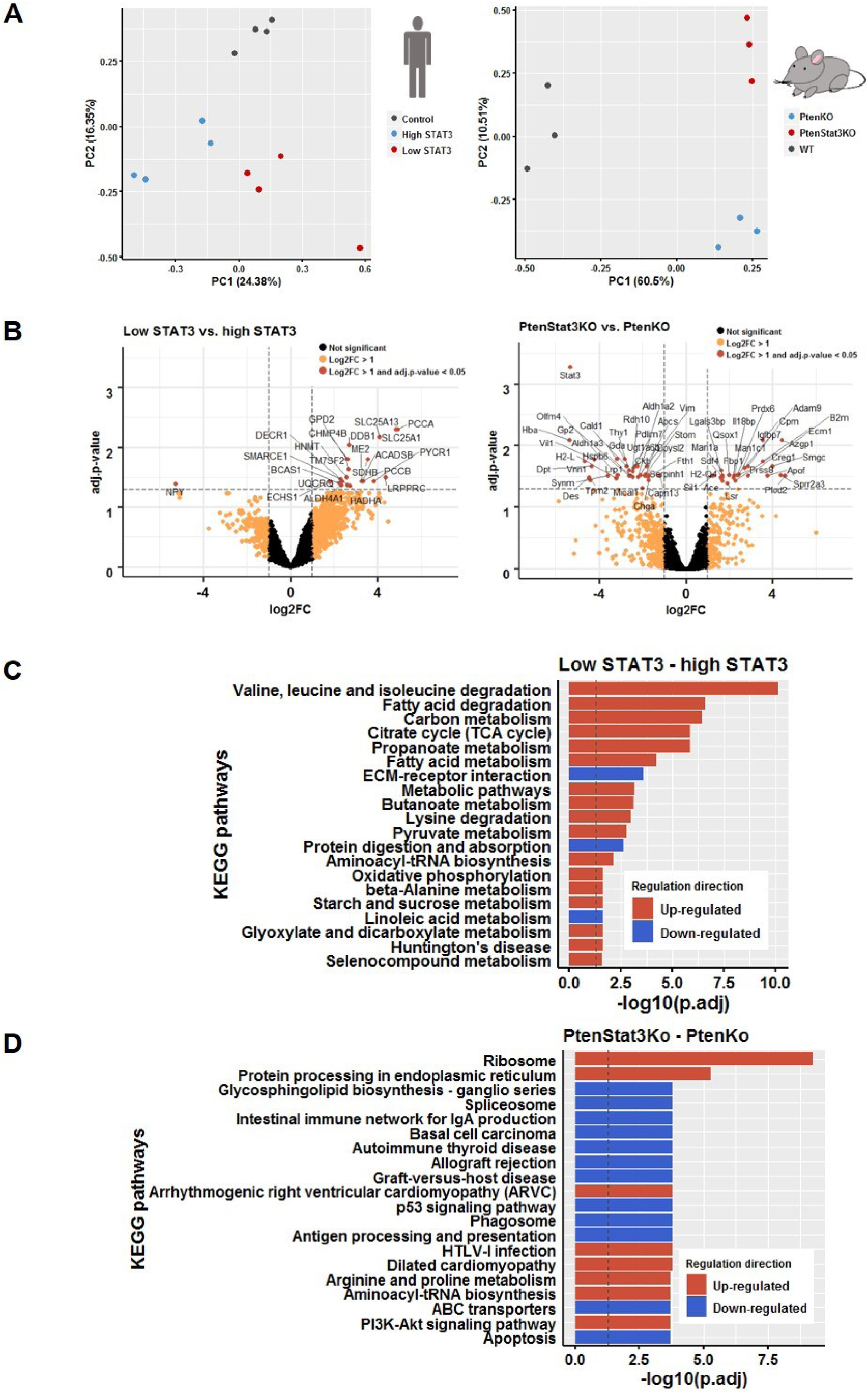
Proteomics from human and murine FFPE-samples show STAT3-dependent profiles. A. PCA of human (left) and murine (right) proteomic samples. Colors represent groups (Human: red = low STAT3, blue = high STAT3, grey = Control; Mouse: red (PtenStat3Ko) = *PtenStat3^pc-/-^*, blue (PtenKo) = *Pten^pc-/-^*, grey = WT). PCA = Principal component analysis. B. Volcano plot of DE proteins of human (left) and murine (right) proteomic samples. Comparison of low STAT3 with high STAT3 (human) and of *PtenStat3^pc-/-^* with *Pten^pc-/-^* (mouse). X-axis represents log2-FC and y-axis -log10 adj. p-values. Colors indicate adj.p-value and log2-FC. (Black = Log2- FC ≤ 1 and adj. p-value ≥ 0.05, orange = Log2-FC > 1 and adj. p-value ≥ 0.05, orange-red = Log2- FC > 1 and adj. p-value < 0.05). Labels indicate gene names of respective proteins. DE = differentially expressed, FC = fold change. See also Tables S3 and S4. C. and D. KEGG pathways enriched in low STAT3 versus high STAT3 (human, C) and *PtenStat3^pc-/-^* versus *Pten^pc-/-^* (mouse, D) groups. PtenKo = *Pten^pc-/-^*, PtenStat3Ko = *PtenStat3^pc-/-^*. See also Tables S3 and S4.

### Proteomics from murine FFPE-samples show increased ribosomal activity in *PtenStat3^pc-/-^-* tumors

We wanted to know if proteomics from a PCa mouse model would reflect the results we obtained from human data. We used a previously established genetic PCa mouse model (Alonzi et al., 2001, Pencik et al., 2015, Suzuki et al., 2001, Wu et al., 2001) (Methods) with conditional loss of either *Pten* (referred to as *Pten^pc-/-^*), or concomitant loss of *Pten* and *Stat3* (*PtenStat3^pc-/-^*) in the prostate epithelium (*Pb-Cre4 Pten^fl/fl^*). Whereas *Pten^pc-/-^* mice show slow, localized tumor progression, the additional deletion of *Stat3* leads to rapid tumor growth, dissemination and early death (Pencik et al., 2015). We selected triplicates from each genotype (wild type (WT), *Pten^pc-/-^* and *PtenStat3^pc-/-^*) and performed LMD and LFQ shotgun proteomics on FFPE tumors and controls (Methods). We were able to detect 2994 proteins on average (2052 - 3465), with 1510 being differentially expressed between all three groups (log-FC ≥ 1, adj. p-value < 0.05). PCA showed a clear separation between groups, and STAT3 was the strongest differentially expressed protein in *PtenStat3^pc-/-^*-compared to *Pten^pc-/-^* tumors (log-FC = -5.34427, adj.p-value = 0.0005; Figure 4a-b Table S4). Comparing *PtenStat3^pc-/-^*- to *Pten^pc-/-^* mice, we found significant upregulation of KEGG pathways associated with ribosome and protein processing in ER. In addition, PI3K-Akt signaling was upregulated, whereas several pathways related to immune response were downregulated (Figure 4D). Gene set testing on GO BP terms, comparing *PtenStat3^pc-/-^*- to *Pten^pc-/-^* tumors, showed upregulation of ribosome biogenesis, translational initiation, rRNA metabolic process, protein localization to ER and establishment of protein localization to ER, among others (Table S4). STAT3-associated regulation of ribosomal activity on protein level was consistent with our human TCGA samples and corresponded to gene cluster 3. In summary, our experimental results indicate a STAT3-dependent repression of ribosomal biogenesis and translation, which suggests that low expression of STAT3 or *Stat3* loss is associated with more aggressive tumors with a high need of energy supply for growth, dissemination and metastasis (Donati et al., 2012).

For comparison with our human proteomic data, we performed gene set testing with a subset of metabolic KEGG pathways, including TCA/OXPHOS, which were shown to be enriched in human proteomic samples (Table S3). We found the TCA cycle to be also significantly upregulated in *PtenStat3^pc-/-^* mouse tumors (Table S4). Since differences in log-FCs were not high between groups, we sought to additionally investigatie *Stat3*-dependent changes in the TCA cycle activity on metabolite level.

### Metabolomics show increased TCA cycle activity in *PtenStat3^pc-/-^* mouse tumors

In order to assess *Stat3*-dependent changes in TCA cycle metabolite levels, we performed a targeted metabolomics experiment on WT, *Pten^pc-/-^* and *PtenStat3^pc-/-^* mice (with biological replicates n= 5 for WT and *Pten^pc-/-^* and n= 3 for *PtenStat3^pc-/-^*, see Methods). Hence, we measured absolute amounts (nmol/µg) of pyruvate, citrate, α-ketoglutarate, succinate, fumarate and malate in mouse prostate tumors and WT prostates. *PtenStat3^pc-/-^* prostate tumors showed significantly higher amounts of pyruvate (Anova with TukeyHSD, adj. p-value = 0.01), fumarate (adj. p-value = 0.027) and malate (adj. p-value = 0.029) compared to WT tumors (Figure 5A, Table S5). Citrate levels were not statistically different between groups (Table S5). Succinate was the only metabolite with lower amounts in *PtenStat3^pc-/-^*- or WT- compared to *Pten^pc-/-^* prostates. Whereas there was a trend of upregulation of TCA cycle metabolites between WT and *Pten^pc-/-^*, only succinate showed a significant difference (adj. p-value = 0.023). Generally, measured metabolite concentrations showed a trend to be higher in *PtenStat3^pc-/-^* compared to *Pten^pc-/-^* mice, but due to the high variability in metabolite levels of *Pten^pc-/-^* mice, significance was not reached in these samples. The results of all ANOVAS with Tukey multiple comparisons of means can be found in the supplementary data (Table S5).

**Figure 5:**
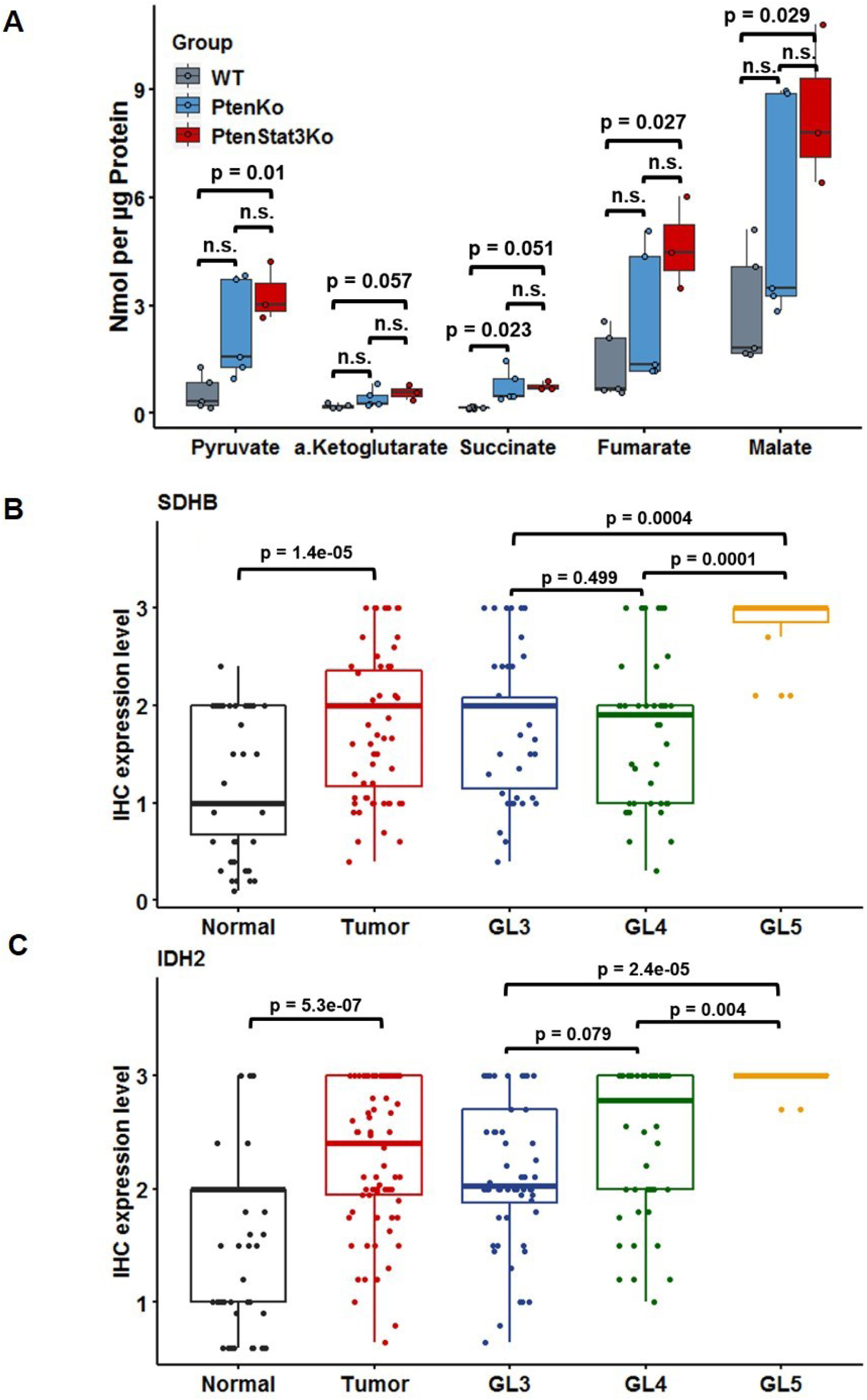
Increased TCA/OXPHOS is associated with tumor aggressiveness in PCa. A. Boxplot representing metabolite concentrations in nmol/µg of 5 metabolites in WT, *Pten^pc-/-^* and *PtenStat3^pc-/-^* prostates. Jitter represents biological replicates. ANOVA test and Tukey multiple comparisons were performed to assess significance. N.s. values are not stated due to better readability, but all p-values can be found in Table S5. Colors represent groups (Red (PtenStat3Ko) = *PtenStat3^pc-/-^*, blue (PtenKo) = *Pten^pc-/-^*, grey (WT) = wild type). N.s. = not significant. X-axis indicates the respective metabolites, y-axis the metabolite concentration in nmol/µg. B. and C. Boxplots representing SDHB (B) and IDH2 (C) protein expression levels detected by IHC. Jitter represents single values in groups. Kruskal-Wallis test and Dunn’s all pairs test were performed to assess significance. GL = Gleason grade, IHC = Immunohistochemistry.

### Increased TCA/OXPHOS is associated with tumor aggressiveness in PCa

By reason of our results of the proteomic analyses, we re-analyzed our TCGA RNA-Seq data for TCA/OXPHOS pathways. We used a subset of metabolic KEGG pathways that was significantly enriched in human proteomic samples for gene set testing of low STAT3 vs. high STAT3 groups. Besides OXPHOS, we also found the TCA cycle to be significantly enriched (Table S1).

It is established, that PCa tumorigenesis is accompanied by enhanced TCA/OXPHOS activity (Costello and Franklin, 2006, Costello et al., 1997, Cutruzzolà et al., 2017). Our data suggest a STAT3-dependent down-regulation of TCA/OXPHOS, supporting the notion of a tumor suppressive function of STAT3 in primary PCa.

To compare TCA/OXPHOS enzyme levels between PCa and healthy prostate, we performed immuno-histochemical (IHC) stainings of a tissue microarray (TMA) consisting of primary PCa and adjacent tumor free tissue from 83 patients (Methods). We stained for SDHB and IDH2. Together with SDHA, SDHC and SDHD, SDHB forms the succinate dehydrogenase (SDH) complex (or respiratory complex II, CII), which is located in the inner mitochondrial membrane. SDH/CII participates in both the TCA cycle by oxidizing succinate to fumarate and OXPHOS by shuttling electrons. IDH2 is the TCA cycle enzyme that converts isocitrate to α-ketoglutarate (Anderson et al., 2018, Stelzer et al., 2016). Both SDHB and IDH2 showed higher expression levels in tumors than in normal tissue (Kruskal Wallis test and Dunn’s all pairs test: SDHB: adj. p-value = 1.4e-05, IDH2; adj. p-value. = 5.3e-07). Moreover, GL5 areas showed a stronger expression of both SDHB (adj. p-value = 0.00044 to GL3 and 0.00014 to GL4) and IDH2 (adj. p-value = 2.4e-05 to GL3 and 0.00461 to GL4), when compared to GL 3 or 4 areas (Figure 5B-C). These data confirm that increased TCA/OXPHOS is associated with tumor aggressiveness in PCa.

### Low-*PDK4* expression is significantly associated with earlier disease recurrence in PCa

Considering that low *STAT3* expression correlates with increased TCA cycle activity and enhanced OXPHOS, we were looking for differentially expressed genes that might cause this effect. Pyruvate dehydrogenase kinase 4 (*PDK4*) was significantly downregulated in low STAT3 RNA-Seq samples (log-FC = -1.126, adj. p-value = 1.47E-07); it is known to inhibit metabolic flux through the TCA cycle and thereby downregulate OXPHOS (Jeoung, 2015; Zhang et al., 2014) (Figure 6A).

**Figure 6:**
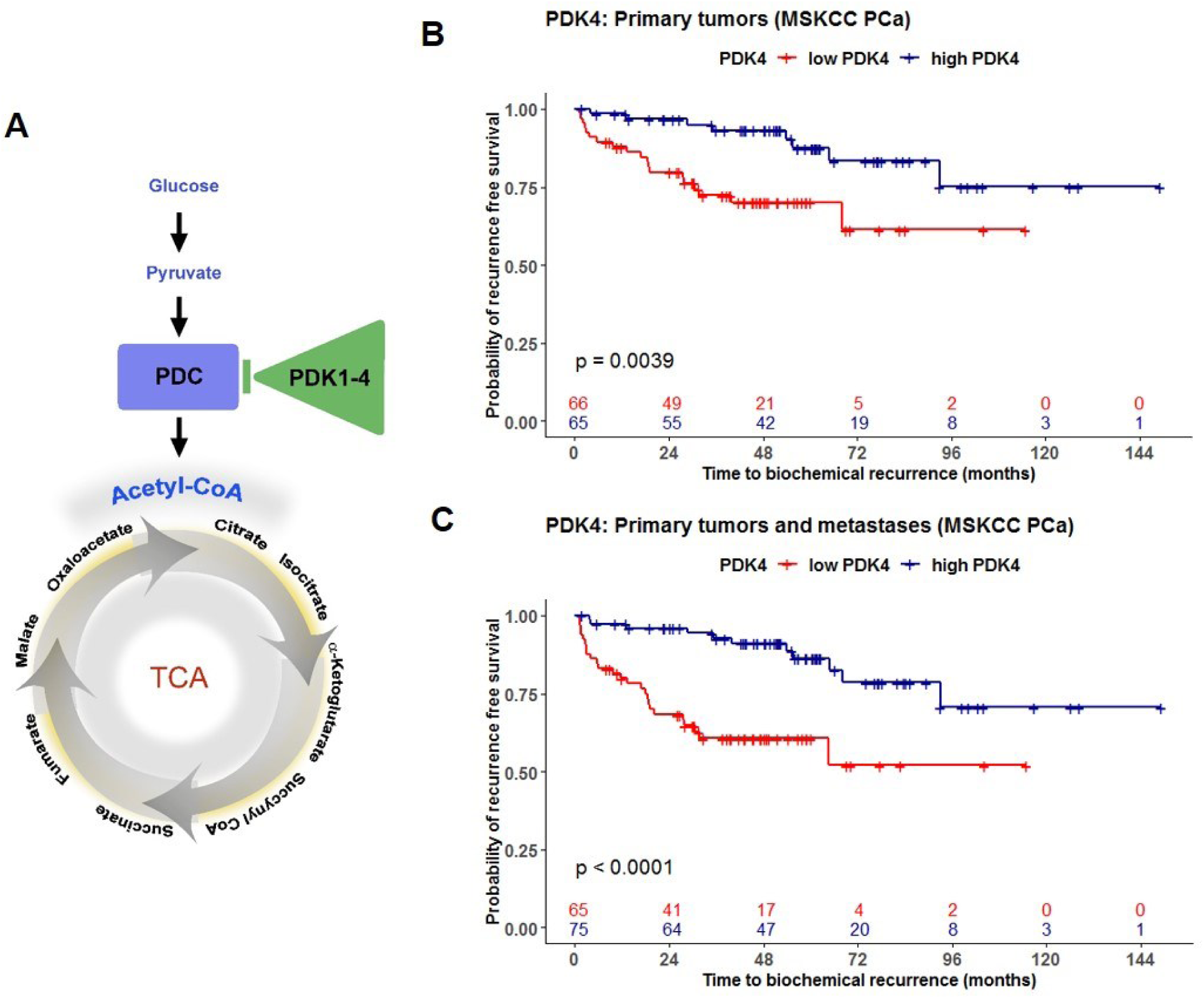
Low *PDK4* is significantly associated with earlier disease recurrence in PCa. A. Simplified scheme of upstream regulation of the TCA cycle. Arrows indicate activation and bar indicates repression. TCA = Tricarboxylic acid cycle, PDC = Pyruvate dehydrogenase complex, PDK = Pyruvate dehydrogenase kinase. B. and C. Kaplan-Meier plots showing time to BCR in months for *PDK4* in primary tumors (B) and in primary and metastatic tumors combined (C) in the MSKCC PCa GSE21032 dataset. Groups were generated by a median split. P-values were estimated by a Log-rank rest. Red = Low *PDK4* expression, blue = High *PDK4* expression, + = censored; See also Figures 7, S3, S4 and S5.

We analyzed the association of *PDK4* expression with BCR in a public gene expression dataset (MSKCC PCa, GSE2103) (Taylor et al., 2010), consisting of 181 primary and 37 metastatic clinically annotated PCa samples. *PDK4* was a significant predictor of BCR both in primary tumors (univariate Cox proportional hazards model: beta: -0.7582, Hazard ratio (HR): 0.4685, p.-value: 0.0011, Figure 6B) and in primary and metastatic tumors combined (beta: -0.9815, HR: 0.3747, p.-value: 1.87e-06, Figure 6C). When compared to diagnostic risk factors, it predicted BCR in low/intermediate risk primary tumors (= clinical staging T1c-T2c) independent of ISUP grades (multivariate Cox proportional hazards model, Figure 7A). In addition, *PDK4* was a significant predictor independent of ISUP grading and clinical tumor staging, as well as pathological tumor staging and pre-surgical PSA-levels in primary and metastatic tumors combined (Figure 7B-C). *PDK4* expression was also significant relative to the occurrence of chemotherapy, hormone therapy and radiation therapy (Figure 7D).

**Figure 7:**
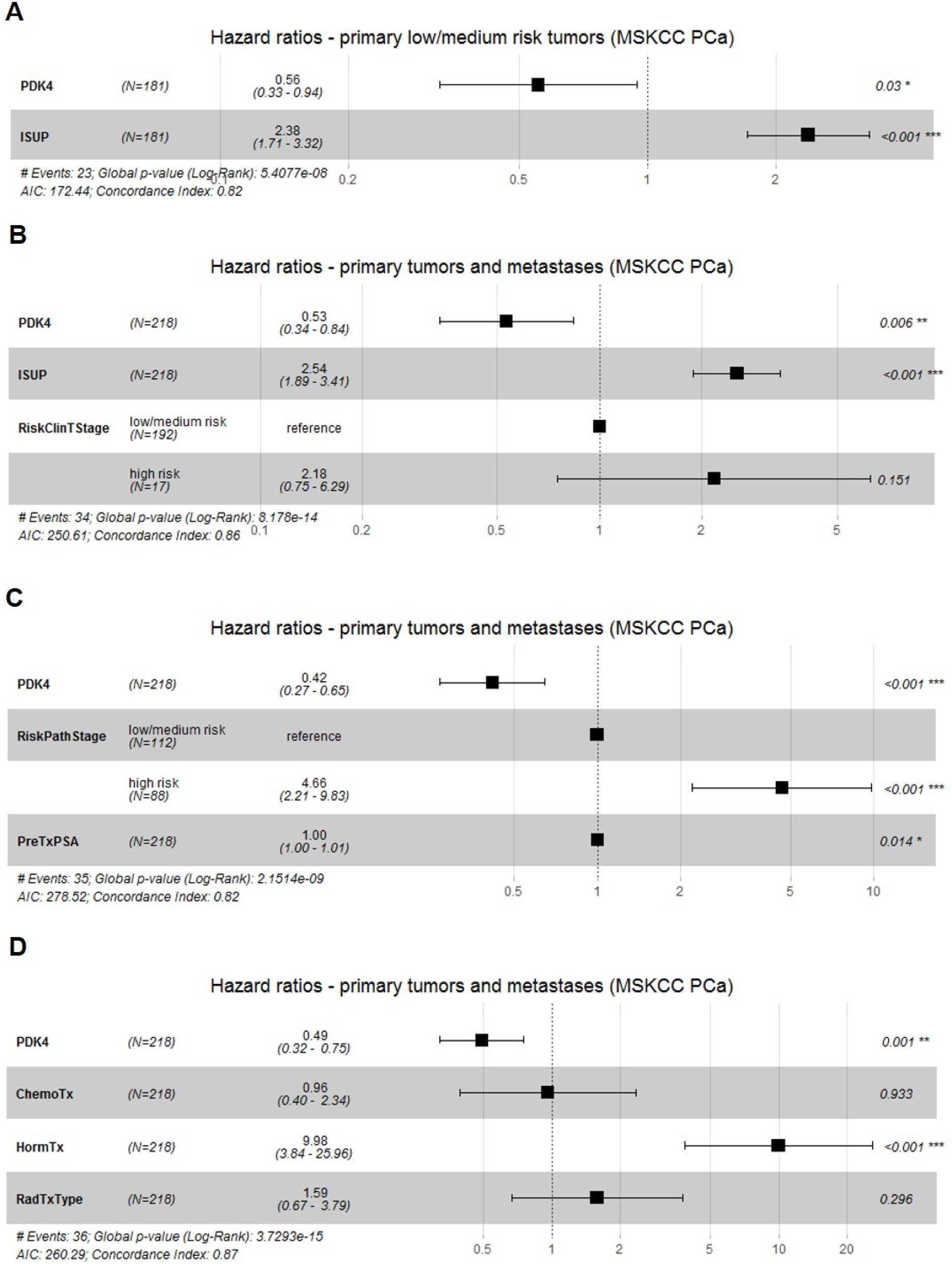
*PDK4* is an independent predictor of biochemical recurrence. A. and B. and C. and D. Forest plots showing hazard ratios for *PDK4* and PCa risk factors in primary low/intermediate risk (T1c-T2c) tumors (A) and in primary and metastatic tumors combined (B-D). Shown from left to right: Name of risk factor, number of samples, hazard ratio (confidence intervals), p-values. ISUP = histological grading from I-V by International Society of Urological Pathology (ISUP) modified Gleason Score, RiskClinTStage = clinical tumor staging (low/intermediate risk = T1c-T2c, high risk = T3-T4), RiskPathStage = pathological tumor staging (low/intermediate risk = pT2a-pT2c, high risk = pT3a-pT4), PreTxPSA = PSA-level prior to prostatectomy, ChemoTx = patient received chemotherapy (no/yes), HormTx = patient received hormone therapy (no/yes), RadTxType = patient received radiation therapy (no/yes).

Considering the possibility of a dataset-specific effect of *PDK4* expression, we additionally tested 4 other datasets with the survExpress tool (Methods) (Aguirre-Gamboa et al., 2013). They all showed a similar trend: low-*PDK4* patients have a higher chance for earlier BCR or death. The Sboner Rubin Prostate dataset (Sboner et al., 2010) consists of survival data of 281 patients with primary PCa from a watchful waiting cohort with up to 30 years clinical follow up. In this cohort, patients in the low-*PDK4*/high-risk group, had a higher chance of earlier death (Risk Groups HR = 1.4 (confidence interval (CI) 1 ∼ 1.98), p-value = 0.05, Figure S3A). Difference in survival for *PDK4* groups was significant for Gleason 6 (Risk Groups HR = 2.92 (CI 1.16 ∼ 7.36), p-value = 0.023, Figure S3B) and 8 (Risk Groups HR = 3.06 (CI 1.06 ∼ 8.85), p-value = 0.039, Figure S3D) and has a p-value = 0.057 in Gleason 7 (Risk Groups HR = 1.85, (CI 0.98 ∼ 3.51), Figure S3C). It has to be considered, that due to the advanced age of PCa patients (85% of all cases are diagnosed in patients >65 years (National Collaborating Centre for Cancer, 2014)) and the duration of the follow-up period of up to 30 years, patients in this study might possibly have suffered from multiple co-morbidities, which also affect survival time.

We also analyzed the TCGA-PRAD dataset (The Cancer Genome Atlas Research Network, 2015) which includes data on survival time, but no reliable information on BCR. Since the overall number of patient deaths is only 10 in this dataset, statistical significance was not reached (HR = 3.14, (CI 0.65 ∼ 15.11), p-value = 0.15). Nevertheless, 8 out of 10 patients who died are in the low-*PDK4*/high-risk group after a median split of sample groups (Figure S4A). We tested two additional datasets – Gulzar (Gulzar et al., 2013) and Lapointe (Lapointe et al., 2004) – that are considerably smaller (n= 89 with 24 events in Gulzar and n = 29 with 7 events in Lapointe). In those, *PDK4* did not reach significance, presumably because of the smaller sample sizes. However, there is a clear trend of low *PDK*4 showing risk of earlier BCR (Figures S4B-C). In conclusion, all datasets revealed a trend of low-*PDK4* patients having a higher risk for earlier BCR or death.

Other tested candidates that were linked to TCA/OXPHOS or regulated ribosomal pathways, such as hypoxia-inducible factor-1α (*HIF-1α*) and CCR4-NOT transcription complex subunit 1 (*CNOT1*), were not predictive of increased risk to earlier disease recurrence (Figure S5A and S5B).

## Discussion

In this study we have used a gene co-expression network analysis in addition to proteomics from laser microdissected human and murine FFPE samples to identify PDK4 as a highly relevant independent candidate prognostic marker in PCa. We here for the first time demonstrate that PCa patients with low *PDK4* expression have a higher risk of earlier disease recurrence, independent of ISUP grading and tumor staging. Moreover, specifically in low-intermediate risk T1c-T2c tumors, *PDK4* proves to be a significant predictor of earlier BCR in the dataset tested, independent of ISUP grading. Therefore, *PDK4* is a strong candidate marker for risk stratification of the large group of T1c-T2c tumors, which are prone to over- or undertreatment. The influence of PDK4 on the PCa disease course may also have an impact on the treatment of type 2 diabetes mellitus (T2DM) with drugs targeting PDK4 as discussed below.

Due to the slow clinical progression rate of PCa, BCR is generally used for risk determination. On account of the protracted nature of a prospective study we evaluated the effect of PDK4 on PCa BCR retrospectively. We therefore believe that it might be beneficial to conduct additional prospective studies to support the postulated effects of PDK4 on PCa outcome.

The generation of mitochondrial adenosine triphosphate (ATP) through aerobic respiration via TCA/OXPHOS is the primary source of energy in most normal cells (Hanahan and Weinberg, 2011, Stacpoole, 2017). Prostate epithelial cells, however, are characterized by a physiological downregulation of TCA/OXPHOS, caused by citrate secretion and zinc accumulation in the cell (Costello and Franklin, 2006, Costello et al., 1997, Cutruzzolà et al., 2017). This is due to the highly specialized role of prostate epithelial cells, which excrete citrate-rich prostatic fluid (Costello et al., 1997). In most cancers, malignant transformation is accompanied by a shift from aerobic respiration via TCA/OXPHOS to aerobic glycolysis, an event also known as the Warburg effect (Hanahan and Weinberg, 2011, Stacpoole, 2017). Primary PCa cells, however, do not show the Warburg effect. On the contrary, the malignant shift from healthy prostate cells to primary PCa cells involves the upregulation of TCA/OXPHOS (Cutruzzolà et al., 2017, Costello and Franklin, 2006). PDK4, which is part of the pyruvate dehydrogenase kinase (PDK) family, plays a central role in the regulation of TCA/OXPHOS (Zhang et al., 2014). PDK4 phosphorylates the pyruvate dehydrogenase complex (PDC) subunits and thereby inhibits the formation of acetyl-coenzyme A from pyruvate. This leads to a down-regulation of metabolic flux through the TCA cycle (Zhang et al., 2014, Jeoung, 2015, Stacpoole, 2017). *PDK4* is a known STAT5 target gene (White et al., 2007) and a putative STAT3 target gene: ENCODE ChIP-Seq against STAT3 in human HeLa-S3 cells shows binding to the promoter region of *PDK4* (Chen et al., 2013, Davis et al., 2018, Kuleshov et al., 2016, The Gene Ontology Consortium, 2018). Thus, the gene expression profile of low STAT3 PCa samples, showing *PDK4* downregulation and TCA/OXPHOS upregulation, reflects a PCa-specific metabolic setting and emphasizes the increased aggressiveness of those cancers compared to high STAT3 tumors.

The crucial role of PDK4 as inhibitor of PDC activity renders it important as a target gene in many cancers and metabolic disorders (Jeoung, 2015, Yamane et al., 2014, Zhang et al., 2014). In non-prostate cancer cells, high *PDK4* facilitates the transition from OXPHOS to aerobic glycolysis and is therefore considered a risk factor enhancing the Warburg effect (Zhang et al., 2014). High *PDK4* is associated with poor survival in breast cancer (Guda et al., 2018) and increased cell growth in bladder cancer cell lines (Woolbright et al., 2018). In addition to direct interaction with PDC, PDK4 has been shown to enhance the Warburg effect via mammalian target of rapamycin (mTOR) and HIF-1α. In mouse embryonic fibroblasts (MEFs) and Eker leiomyoma tumor-3 (ELT3) cells, Liu et al. show, that PDK4 activates mTOR signaling via cAMP-response element-binding Protein (CREB) and Ras homolog enriched in brain (RHEB) (Liu et al., 2014). The mTOR effector HIF-1α and its downstream target pyruvate kinase isozyme M2 (PKM2) were elevated in *PDK4* overexpressing cells and reduced in *PDK4* knockdown cells. Both HIF-1α and PKM2 have been known to modulate key processes required for the Warburg effect (Courtnay et al., 2015).

Conversely, in cancer cells that have undergone tumor progression via epithelial-mesenchymal transition (EMT), a low-*PDK4*-mediated metabolic shift from glycolysis to OXPHOS was reported, and knockdown of *PDK4* was sufficient to induce EMT in human non-small cell lung cancer (NSCLC) cell lines (Sun et al., 2014). In accordance with these findings, Sun et al. show reduced overall survival of NSCLC patients with low *PDK4* expression. Yang et al. show that downregulation of PDK4 is associated with earlier recurrence and lower survival time in hepatocellular carcinoma (Yang et al., 2019). In concurrence with our data, Chen et al. (Chen et al., 2018) show that prostate tumors exhibit higher gene expression and higher protein levels of both PDC subunit pyruvate dehydrogenase A1 (PDHA1) and the PDC activator pyruvate dehydrogenase phosphatase 1 (PDP1). Mengual et al. find PDK4 to be significantly higher expressed in both tumors and post-prostatic massage urine samples from PCa patients compared to the respective control groups (Mengual et al., 2014).

*PDK4* is elevated in patients with T2DM and other metabolic disorders involving insulin resistance, such as obesity (Kulkarni et al., 2012, Lee, 2014, Stacpoole, 2017). Furthermore, it has been established that the development of diabetes and insulin resistant states is causally linked to PDC inhibition through PDK upregulation (Stacpoole, 2017, Jeoung, 2015). Therefore, the PDC/PDK4 axis is an important therapeutic target in the treatment of both diabetes and cancer. Dichloroacetate (DCA), for example, is a PDK inhibitor that is most active against PDK2, but also against PDK1 and PDK4, and has been used as an investigational drug for over 30 years in diabetes, cancer and other diseases (Stacpoole, 2017). It has been subjected to a large number of clinical trials but has lacked pharmaceutical support due to its non-patentability. Attempts to synthesize similar small-molecule inhibitors of PDK were made, but failed in clinical trials (Stacpoole, 2017). Nonetheless, PDK inhibitors remain promising targets in the treatment of T2DM and cancer.

T2DM is reported to have a protective effect on the development of PCa (Baradaran et al., 2009, Choi et al., 2016). Our indication of a beneficial effect of PDK4 upregulation on the clinical course of PCa suggests that a T2DM-induced inhibition of the TCA cycle via the PDC/PDK4-axis is a potential cause of this protective effect, thereby leading to an indolent disease or potentially preventing the development of high risk PCa. As a note of caution, our findings suggest that therapeutic targeting of PDK4 in patients with both T2DM and PCa may result in increased tumor aggressiveness. Hence, our data may be of high clinical importance.

For proteomic characterization, we used human FFPE-material up to 21 years old. Patient material is routinely processed and stored as FFPE specimen for diagnostic purposes in the clinic. Academic pathological institutions possess large archives of fully annotated FFPE-patient material that can be used for retrospective research. Recent developments in LC-MS/MS sample preparation protocols have made it possible to use those archives for proteomic sample-characterization (Ostasiewicz et al., 2010, Wisniewski et al., 2013). This is particularly useful in diseases such as PCa, that frequently have a protracted clinical progression rate and may take years for disease recurrence and development to metastatic disease to occur. Previous proteomic studies focused on the characterization of the proteome (and transcriptome) of primary prostate cancer (Iglesias-Gato et al., 2016, Sinha et al., 2019), on proteomic and transcriptomic disease evolution (Latonen et al., 2018) or on the establishment of a diagnostic panel via machine learning (Kim et al., 2016). In contrast to the above studies, our approach is focused on the specific effects of loss of the key transcription factor STAT3 in prostate cells only.

Although STAT3-signaling is linked to various regulatory events causing increased proliferation, stemness and inflammation and therefore has oncogenic properties, STAT3 can also act as tumor suppressor (Huynh et al., 2019). The deletion of *Stat3* in prostate epithelial cells in a loss of *Pten* PCa mouse model leads to increased tumor growth and early death (Pencik et al., 2015). It is well established that STAT3 is able to control the activity of mitochondria, the electron transport chain (ETC) and the ER both via transcriptional control and via direct binding to these cell compounds (Avalle et al., 2018, Huynh et al., 2019, Poli and Camporeale, 2015, Wegrzyn et al., 2009). On the one hand, *STAT3* expression is associated with increase in glycolysis and the suppression of the ETC (Wegrzyn et al., 2009, Huynh et al., 2019). Specifically, the activation of STAT3 is linked to the induction of HIF-1α, which suppresses OXPHOS and reprograms TCA (Camporeale et al., 2014, Demaria et al., 2010, Niu et al., 2008, Pawlus et al., 2014, Poli and Camporeale, 2015). Likewise, HIF-1 was shown to transcriptionally upregulate PDK4 (Courtnay et al., 2015). On the other hand, however, STAT3 can be directly associated with mitochondrial complexes, improving ETC activity and transcription of mitochondrial genes (Huynh et al., 2019, Wegrzyn et al., 2009). Our data support the former concept of STAT3-dependent downregulation of TCA/OXPHOS in accordance with HIF-1α upregulation and additionally suggest inhibition of the TCA cycle via PDK4. We did not find a significant influence of *HIF-1α* on the relapse of PCa, nor was *CNOT1*, which is associated with inhibition of ribosomal translation initiation, significantly associated with a worse survival outcome.

In summary, the present study uses a systems-biology approach to unveil the effects of loss of STAT3 in PCa. In this setting we show an association of STAT3 to TCA/OXPHOS, ribosomal biogenesis and translation. Our data do not only substantiate previous research on the effects of STAT3, but in a prostate specific context also explain the tumor suppressive functions of STAT3 from a cell autonomous point of view. We here identify *PDK4* as a promising independent prognostic marker for PCa which will facilitate to separate good from bad prognostic PCa. In addition, the known protective effect of diabetes on the development of PCa could be owing to the upregulation of PDK4. Since PDK4 inhibitors are promising therapeutics in T2DM, these drugs may potentially increase the aggressiveness of PCa. Therefore, our results are of high general and clinical importance, and further studies on the function of PDK4 in PCa are urgently needed.

## Supporting information

Table S1 related to Figure 1, S1 and S2

Table S2, related to Figure 2 and 3

Table S3 related to Figure 4

Table S4 related to Figure 4

## Acknowledgments

We thank Kathrin Oberhuber and Karin Nowikovsky for editing the manuscript. We thank Prof. Christoph Herwig (Institute of Chemical Engineering, Bioprocess Technology, Vienna University of Technology) for access to resources. We thank Saptaswa Dey, Paul Kroll, Daniela Dunkler and Alexandra Kaider for insightful discussion. This work was funded by the COMET Competence Center CBmed - Center for Biomarker Research in Medicine (FA791A0906.FFG). The COMET Competence Center CBmed is funded by the Austrian Federal Ministry for Transport, Innovation and Technology (BMVIT); the Austrian Federal Ministry for Digital and Economic Affairs (BMDW); Land Steiermark (Department 12, Business and Innovation); the Styrian Business Promotion Agency (SFG); and the Vienna Business Agency. The COMET program is executed by the FFG.

## Author contributions

Conceptualization, MO, BH, LK;

Methodology, MO, MP, MR, GK, JG, TM, AHo, GO, GE, BH and LK.

Validation, MO, MP, MR, MW, MSchl and JPe;

Formal Analysis, MO, MP and MR;

Investigation, MO, MP, MR, MW, GO, PH, JPe, RW, EG, AHo, TW, MSchm and MSchl;

Resources, GO, AHa, TM, JPo, GK, MM and LK

Data Curation, MO, MP and MR;

Writing – Original Draft, MO;

Writing – Review & Editing, MO, GO, JG, GE, MB, BH and LK;

Visualization, MO and AJ;

Supervision, LK, BH, GO, JG and TM;

Project Administration, MB, WW, BH and LK;

Funding Acquisition, WW, BH and LK

## Methods

### CONTACT FOR REAGENT AND RESOURCE SHARING

Further information and requests for resources and reagents should be directed to and will be fulfilled by the Lead Contact, Lukas Kenner.

### EXPERIMENTAL MODEL AND SUBJECT DETAILS

#### Clinical specimens

FFPE- prostate material was obtained from the Department of Pathology of the Medical University of Vienna (MUW), Vienna, Austria. The FFPE-material originated from 84 primary PCa patients and 7 bladder cancer (BCa) patients who underwent radical prostatectomy at the General Hospital of Vienna (AKH) from 1993 to 2015. Use of patient FFPE-material in this study was approved by the Research Ethics Committee of the Medical University Vienna, Austria (1877/2016).

#### Animal model

Mice carrying a prostate specific deletion of *Pten* (*Pten^pc-/-^*) were received from Prof. Johannes Schmidt (Birbach et al., 2011). They were generated by crossing *Pten^tm2Mak^* (*Pten^loxP/loxP^*) mice (Suzuki et al., 2001) with male *PB-Cre4* transgenic mice (RRID:IMSR_NCIMR:01XF5) (Wu et al., 2001). Furthermore, mice carrying *Stat3 ^loxP/loxP^* (Alonzi et al., 2001) were crossed with *Pten^pc-/-^* mice to obtain mice with a concomitant loss of *Pten* and *Stat3* (*PtenStat3^pc-/-^*) in the prostate epithelium (Pencik et al., 2015). All mice were maintained on a C57BL/6 and Sv/129 mixed genetic background. Animal experiments were reviewed and approved by the Austrian ministry authorities and conducted according to relevant regulatory standards (BMWFW-66.009/0281-I/3b/2012 and BMWFW-66.009/0088-WF/V/3b/2018). Mice were housed on a 12- 12 light cycle (light on 6 am and off 6 pm) and provided food and water ad libitum. For experiments, 19 week old male mice were used. All efforts were made to minimize suffering.

### METHOD DETAILS

#### TCGA-PRAD RNA-Seq data acquisition

TCGA PRAD (The Cancer Genome Atlas Research Network, 2015) RNA-Seq data were acquired as HTSeq-Counts from GDC Legacy archive via R package TCGAbiolinks v.2.10.5 (Colaprico et al., 2016). Only primary tumor samples (n=489) were selected. For data pre-processing, R package edgeR v.3.24.3 (Robinson et al., 2010) was used. Raw data were transformed to counts per million (cpm) values and genes that were expressed in less than 70% of samples were omitted. Gene expression distributions were normalized using weighted trimmed mean of M-values (TMM) method (Robinson and Oshlack, 2010). Samples were ranked according to *STAT3* expression and assigned to groups: “high STAT3” consisted of the 1- 0.8^th^ quantile (n=100), “low STAT3” of the 0.2^nd^ quantile (n=100) and “medium STAT3” of all samples in between (n=298).

#### Weighted gene co-expression network analysis (WGCNA)

TCGA PRAD RNA-Seq data were used to generate a weighted gene co-expression network with WGCNA v.1.66 R package as described by Langfelder and Horvath (Langfelder and Horvath, 2008, Langfelder and Horvath, 2012). For creation of a trait matrix, TCGA PRAD clinical data were acquired via GDC Legacy Archive. Patients without information on disease recurrence were excluded. Following clinical traits were used for analyses: Biochemical disease recurrence (BCR), pathological tumor staging (pT), pathological lymph node staging (pN) and histological grading with Gleason Score (GSC). Pathological staging was split into low to intermediate risk (indicated as 1) and high to very high risk (indicated as 2) groups. For pT, the low to intermediate risk group consisted of T2abc- and the high to very high risk group of T3-T4 samples. For pN, low to intermediate risk was assigned to N0 samples, high to very high risk to N1 samples. The emergence of BCR was indicated as 1, otherwise as 0. GSCs were not split into groups. *STAT3*-expression was included from RNA-Seq data.

RNA-Seq data was acquired and prepared as described above. Only samples with matching trait data were used for network creation (n=397). Gene expression data was voom-transformed with limma v.3.38.3 R package (Ritchie et al., 2015, Law et al., 2014) and outliers were removed by sample clustering. 382 samples and 13932 genes were used for network construction.

First, a correlation matrix was created using biweight midcorrelation of genes. Second, an adjacency matrix was established from the correlation matrix with a soft thresholding power beta of 6. Third, a topological overlap matrix (TOM) was calculated from the adjacency matrix (Zhang and Horvath, 2005). The TOM provides information on the interconnectedness of genes by a similarity measure: it indicates, whether two genes share co-expression to a similar set of other genes (Zhang and Horvath, 2005, Yip and Horvath, 2007). For the creation of gene clusters (= modules), hierarchical clustering based on TOM-based dissimilarity was performed. Minimum gene cluster size was set to 30. Genes which did not belong to any cluster were summarized as cluster 13. To compare expression profiles of gene clusters, the 1^st^ principal component (= module eigengene (ME)) of each cluster was calculated and clusters with similar eigengenes (r > 0.75) were merged. Genes in each gene cluster were tested for overrepresentation of GOs and KEGG pathways with clusterProfiler v.3.10.1 (Yu et al., 2012). GOs CC, Molecular Function (MF) and BP were tested separately. Significance was defined by an adj. p-value ≤ 0.05, adjustment method was Benjamini-Hochberg.

Gene clusters were associated to external traits by correlating MEs with trait data (= cluster-trait correlation) by Pearson correlation. Student asymptotic p-values for given correlations were adjusted by Benjamini-Hochberg method. Likewise, correlation of each gene to both the respective gene cluster (= module membership, MM) and *STAT3*-expression (= Gene significance, GS) was calculated by Pearson correlation. Student asymptotic p-values were calculated and adjusted with Benjamini-Hochberg method. Significance was defined by an adj. p-value ≤ 0.05.

We defined a strong correlation to be between ±0.6 - ±1, a moderate correlation to be between ±0.59 - ±0.3 and a weak/no correlation between ±0.29 - 0. Two clusters were strongly negatively correlated to STAT3-expression (ρ ≤ - 0.6, adj. p-value ≤ 0.01). For both clusters, genes were sorted for their MM and GS. The top 50 genes with a MM ≥ 0.8 and a GS ≤ -0.6 (adj.p-value ≤ 0.05) were used for overexpression analysis with clusterProfiler (Yu et al., 2012). GO BP enrichment was additionally performed using Cytoscape v.3.6.1. (Shannon et al., 2003) and the ClueGO plug-in v.2.5.1 (Bindea et al., 2009) on those genes.

#### Human tissue micro array (TMA) generation

For generation of a TMA, we used FFPE-material from a patient cohort of 83 patients with primary PCa who underwent radical prostatectomy from 1993 to 2003. The TMA consists of 2 spots from tumor, prostatic intraepithelial neoplasia (PIN) and normal prostate areas from the same patient. Whole mount prostate FFPE-blocks were sliced into 3 µm thick sections, mounted on slides and stained with hematoxylin and eosin. Subsequently, a pathologist marked the respective areas on the slides. To generate the TMA, cores of 2 mm diameter were cut out of the donor block and placed into the recipient TMA block using a manual tissue arrayer (Beecher Instruments). Tissue sections (3 µm thick) were placed onto superfrost slides.

#### Immunohistochemistry (IHC)

IHC was performed on FFPE TMAs using consecutive sections. The following antibodies were used: anti-IDH2 (rabbit polyclonal, 1:100 dilution; Proteintech, 15932-1-AP, RRID: AB_2264612) and anti-SDHB (mouse monoclonal, 1:100 dilution; Abcam; ab14714, RRID: AB_301432). Staining was performed using the Benchmark Ultra automated staining system (Ventana, Roche). The procedure was conducted as follows: heat pre-treatment, antigen retrieval with CC1 buffer (pH 8.5) for 64 min., incubation with the antibody for 32 min. and counterstaining with hematoxylin and bluing reagent for 8 min. each. After automated staining, the slides were washed with water, then dehydrated in increasing concentrations of ethanol (70%, 80%, 96%, absolute alcohol) until xylol, covered with the mounting medium Shandon Consul-Mount^TM^ (Thermo Scientific) and analyzed by standard light microscopy. Antibodies were validated for FFPE IHC. As positive controls, human colon cancer for IDH2 and human muscle tissue for SDHB were used.

#### Sample selection and preparation for laser microdissection (LMD)

From the TMA and patient cohort described above, STAT3 protein expression was quantified by a pathologist after IHC staining (Pencik et al., 2015). We selected 7 patients with no STAT3 expression (0 positive cells) as low STAT3 group and 7 patients with ≥ 11% positive cells as high STAT3 group (see also supplementary information). Additionally, 7 healthy prostate FFPE samples were included as control group, stemming from BCa patients. To facilitate LMD, we created a TMA for each patient. Whole mount prostate FFPE-blocks were sliced into 3 µm thick sections, mounted on slides and stained with hematoxylin and eosin. A pathologist marked tumor areas with GL 4 or 5 on the slides. For each patient, a TMA block was created with 2 mm diameter spots using a manual tissue arrayer (Beecher Instruments).

For LMD of murine samples, FFPE tumor material was used from WT, *Pten^pc-/-^* and *PtenStat3^pc-/-^* mice (n=3 for each genotype). Blocks were sliced into 3 µm thick sections, mounted on slides and stained with hematoxylin and eosin. Tumor areas were marked by a pathologist. Since mouse tumors are much smaller and contain only few stroma compared to human PCa, there was no need to create sample-TMAs for LMD.

#### LMD for proteomic analysis

For LMD of human samples, a Palm Zeiss Microbeam 4 was used. Sample TMA blocks were cut into 10 μm thick sections and mounted on superfrost slides. For LMD of mouse samples, a Leica LMD6000 was used. Tissue blocks were cut into 10 μm thick sections and mounted on membrane slides (PEN Membrane, 2.0 µm, Leica). LMD was conducted similarly for mouse and human samples: for each sample, a slide was stained with hematoxylin and eosin for inspection before LMD. To obtain the minimum amount of tissue (100 nl = 0.1 mm³) necessary for consecutive LC-MS/MS analysis, at least 10 mm² of target area was laser-microdissected. To obtain proteomic profiles solely from the tumor, stroma and immune cells were excluded from dissection. Microdissected FFPE-samples were stored at -20°C before LC-MS/MS analysis.

#### Proteomic liquid chromatography tandem mass spectrometry (LC-MS/MS) measurements

##### Protein extraction and enzymatic digestion

100 nl (10 mm^2^ of 10 µm slides) of FFPE-material per sample was used for analysis. Lysis of microdissected tissue was carried out in 50% trifluoroethanol (TFE), 5 mM dithiothreitol (DTT), 25 mM ammonium bicarbonate (ABC) at 99°C for 45 min. followed by 5 min. sonication (Bioruptor, Diagenode). After centrifugation at 16,000 g for 10 min., the cleared protein lysate was alkylated with 20 mM iodoacetamide for 30 min. at room temperature. Upon vacuum centrifugation, digestion was carried out in 5% TFE, 50 mM ABC to which 0.15 µg of LysC and 0.15 µg of trypsin were added for digestion overnight at 37°C. The following day, digestion was arrested by adding trifluoroacetic acid (TFA) to 1% and the digestion buffer removed by vacuum centrifugation. Peptides were suspended in 2% acetonitrile, 0.1% TFA and purified on C18 StageTips. Finally, purified peptides were resolved in 2% acetonitrile, 0.1% TFA and the entire sample was injected for MS analysis in a single shot measurement. Protocols were adapted from Roulhac et al. and Wang et al. (Wang et al., 2005, Roulhac et al., 2011).

##### LC-MS/MS analysis

LC-MS/MS analysis was performed on an EASY-nLC 1000 system (Thermo Fisher Scientific) coupled on-line to a Q Exactive HF mass spectrometer (Thermo Fisher Scientific) with a nanoelectrospray ion source (Thermo Fisher Scientific). Peptides were loaded in buffer A (0.1% formic acid) into a 50cm long, 75 μm inner diameter column in house packed with ReproSil-Pur C18-AQ 1.9 μm resin (Dr. Maisch HPLC GmbH) and separated over a 270 minute gradient of 2-60% buffer B (80% acetonitrile, 0.1% formic acid) at a 250 nl/min flow rate. The Q Exactive HF operated in a data dependent mode with full MS scans (range 300-1,650 m/z, resolution 60,000 at 200 m/z, maximum injection time 20 ms, AGC target value 3e6) followed by high-energy collisional dissociation (HCD) fragmentation of the five most abundant ions with charge ≥ 2 (isolation window 1.4 m/z, resolution 15,000 at 200 m/z, maximum injection time 120 ms, AGC target value 1e5). Dynamic exclusion was set to 20s to avoid repeated sequencing. Data were acquired with the Xcalibur software (Thermo Scientific).

##### LC-MS/MS data analysis

Xcalibur raw files were processed using the MaxQuant software v.1.5.5.2 (Cox and Mann, 2008), employing the integrated Andromeda search engine (Cox et al., 2011b) to identify peptides and proteins with a false discovery rate (FDR) of < 1%. Searches were performed against the Human or Mouse UniProt database (August 2015), with the enzyme specificity set as “Trypsin/P” and 7 as the minimum length required for peptide identification. N-terminal protein acetylation and methionine oxidation were set as variable modifications, while cysteine carbamidomethylation was set as a fixed modification. Matching between runs was enabled in order to transfer identifications across runs, based on mass and normalized retention times, with a matching time window of 0.7 min. Label-free protein quantification (LFQ) was performed with the MaxLFQ algorithm (Cox et al., 2014, Schaab et al., 2012, Tyanova et al., 2016a, Tyanova et al., 2015, Cox et al., 2011a, Cox and Mann, 2008) where a minimum peptide ratio count of 1 was required for quantification. Data pre-processing was conducted with Perseus software (Tyanova et al., 2016b); v.1.5.8.6 was used for human data and v.1.5.5.5 for mouse data. Data was filtered by removing proteins only identified by site, reverse peptides and potential contaminants. After log2 transformation, biological replicates were grouped and outlier removal was conducted for human samples. For human samples, we pursued analysis with the 4 most similar samples of each group after unsupervised hierarchical clustering. For mouse samples, we continued analyses with three replicates per group. LFQ intensities were filtered for valid values with a minimum of 3 valid values per group, after which missing data points were replaced by imputation. The resulting datasets were exported for further statistical analyses using R. Filtered, normalized and log2 transformed data were imported and PCA and unsupervised hierarchical clustering was performed. Plots were generated with ggplot2 v.3.1.1. (Wickham, 2016.), gplots v.3.0.1.1 (Gregory R. Warnes et al., 2019) and EnhancedVolcano v.1.0.1 (Blighe, 2019) R packages. Differential expression was conducted as described in the “Statistical Information” section on differential expression analysis.

#### Metabolomic liquid chromatography high-resolution mass spectrometry (LC-HRMS) measurements

##### Standards and solvents

Acetonitrile (ACN), methanol (MeOH) and water were of LC-MS grade and ordered at Fisher Scientific (Vienna, Austria) or Sigma Aldrich (Vienna, Austria). Ammonium bicarbonate, ammonium formate and ammonium hydroxide were ordered as the eluent additive for LC-MS at Sigma Aldrich. Formic acid was also of LC-MS grade and ordered at VWR International (Vienna, Austria). Sodium hydroxide (NaOH) was ordered from Sigma Aldrich (Vienna, Austria). Metabolite standards were purchased from Sigma Aldrich (Vienna, Austria) or Carbosynth (Berkshire, UK).

##### Sample preparation of mouse organs

Analysis was conducted with n = 5 biological replicates of wild type and Pten^pc-/-^ mice and n= 3 biological replicates of PtenStat3^pc-/-^ mice. For wild type and Pten^pc-/-^, technical replicates (n=3) were made from 2 biological samples in each group. For PtenStat3^pc-/-^ mice, technical replicates (n=3) were made for all biological samples. The prostates of sacrificed 19 week old mice were immediately collected, quickly washed in fresh PBS, snap-frozen in liquid N_2_, and stored on dry ice in Petri-dishes until extraction. Tissue pieces were transferred into glass vials and 50 µl of fully ^13^C labelled internal standard from ISOtopic solutions e.U. (Vienna, Austria) and 950 µl extraction solvent were added (80% MeOH, 20% H_2_O, both LC-MS-grade (Sigma Aldrich, Vienna, Austria)). Subsequently, the tissue was homogenized with a probe sonicator head (Polytron PT 1200E handheld homogenizer, Kinematica) in the extraction solvent. After homogenization, the contents of the glass tubes were transferred to a 2 ml Eppendorf-tube and the glass tubes were washed two more times with 500 µl extraction solvent to transfer all tissue content to the Eppendorf-tube. The Eppendorf-tubes were thoroughly vortexed and kept on dry ice during the processing of other samples.

The samples were centrifuged (14,000 g, 4°C, 20 min), four 400 µl aliquots were extracted into LC-vials and 3x 100 µl were used for pooled quality controls (QC) for each sample, respectively. Remaining extraction solvent on the pellets was discarded. Aliquots were evaporated until dryness in a vacuum centrifuge. The dried samples and the high molecular pellets were stored at -80°C until measurement.

##### Quantification of metabolites with LC-HRMS

Before the LC-HRMS analysis, the samples were reconstituted in water with thorough vortexing, diluted either 1 to 100 or 1 to 40 in water, and adjusted to be in total of 500 µl 50: 50 H_2_O: ACN.

Quantification was carried out by external calibration using U13C-labelled internal standards. The internal standard always originated from the same aliquot as used for the extraction, and was diluted to the same extent as the sample.

The LC-HRMS measurement was adopted from Schwaiger et al. (Schwaiger et al., 2019). Shortly, a SeQuant® ZIC®-pHILIC column (150 × 2.1 mm, 5 μm, polymer, Merck-Millipore) was utilized with a 15-minute long gradient and 10 mM ammonium bicarbonate pH 9.2/10% ACN and 100% ACN as eluents. Sample measurements were randomized, and within every 10 injections a pooled QC sample, a QC with standards and a blank was injected. HRMS was conducted on a high field Thermo Scientific™ Q Exactive HF™ quadrupole-Orbitrap mass spectrometer equipped with an electrospray source. Full mass scan data with resolution of 120 000, maximum injection time (IT) of 200 ms, automatic gain control (AGC) target of 1e6 in the mass range of 65-900 m/z was acquired with positive-negative-polarity switching.

Targeted analysis of the metabolomics data was carried out with Thermo Trace Finder 4.1 software. In all cases the [M-H]^−^ ion was extracted with 5 ppm mass tolerance.

##### Quantification of total protein content from pellet

The pellets resulting from extraction with 80% MeOH were dissolved in 0.2M NaOH solution, diluted 1:10 and quantified for total protein content with the Micro BCA Protein Assay kit from Thermo (Rockford, USA), according to the manufacturer’s instructions.

Absolute metabolite amounts were normalized to the protein content. If multiple technical replicates were available from the same organ, the sum of metabolite concentrations was calculated for each metabolite and it was normalized with the sum of total protein content for that organ. This way, absolute metabolite amounts (nmol) were normalized to the total protein content (µg) from the tissue of origin.

### QUANTIFICATION AND STATISTICAL ANALYSIS

#### Statistical analysis of WGCNA clusters

Statistical analyses were conducted as described in the section “Weighted gene co-expression network analysis (WGCNA)”. Generally, correlations were assessed by Pearson correlation and Students asymptotic p-values were calculated. P-values were adjusted by Benjamini-Hochberg method. Significance was defined as adj. p-value ≤ 0.05.

#### Human TMA quantification

For statistical evaluation of a human TMAs, IHC stainings on tumor and normal tissue were evaluated by a pathologist. Stainings were quantified by evaluating staining intensities and percentage of positive cells, described by the IHC-expression level (EL):

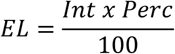

where staining intensity (Int) ranges from 0-3 and percentage of positive cells (Perc) from 0-100. Therefore, ELs can take values from 0-3. To compare GLs, ELs were evaluated separately for each present GL in a spot. To test for significant differences between groups, Kruskal-Wallis test was applied after rejection of null-hypothesis for normality testing with Pearson chi-square normality test. Visual inspection of the distribution of data was conducted using Q–Q (quantile-quantile) plots and density plots. Pairwise comparisons were done using Dunn’s all-pairs test. Significance was defined by adj. p-value ≤ 0.05, adjustment method was Benjamini-Hochberg. Statistical tests were performed using the R software environment with packages DescTools v.0.99.28 (Signorell and al., 2018), PMCMRplus v.1.4.1 (Pohlert, 2018) and nortest v.1.0-4 (Gross and Ligges, 2015). Plots were generated with ggpubr v.0.2 (Kassambara, 2018). Data was processed using tidyverse R packages v.1.2.1 (Wickham, 2017).

#### Survival analysis

Survival analyses were performed using R packages survival v.0.4.5 (Therneau and Grambsch, 2000, Therneau, 2015) and survminer v.2.44-1.1 (Kassambara et al., 2019). Univariate and multivariate Cox proportional hazards models were fitted to assess influence of *PDK4* on BCR. As a rule of thumb, one predictor per 10 events was included in the model. In addition, Kaplan-Meier curves and log-rank tests were performed after a median split of samples by *PDK4*-expression. All statistical tests were considered significant with a p-value ≤ 0.05. BCR is defined by an increase of > 0.2ng/ml PSA in serum on two occasions.

Additional datasets were analyzed using SurvExpress online tool (Aguirre-Gamboa et al., 2013). Here, prognostic index (PI) of tested genes was estimated by fitting a Cox proportional hazards model. Risk groups were generated by ranking samples by their PI (with high PI indicating a high risk) followed by either a median split or a maximizing split (with a split point where the p-value is minimum). Median split was used for *HIF-1α* and *CNOT1* in the MSKCC PCa dataset and for *PDK4* in the TCGA PRAD and Lapointe dataset, maximizing split for the remainder. Risk groups were analyzed by a concurrent Cox model and used for Kaplan-Meier plots and log-rank tests. All statistical tests were considered significant with a p-value ≤ 0.05.

Following publicly available data sets were used for analyses: MSKCC PCa GSE21032 (Taylor et al., 2010), Sboner Rubin Prostate GSE16560 (Sboner et al., 2010), TCGA PRAD (The Cancer Genome Atlas Research Network, 2015), Gulzar Prostate GSE40272 (Gulzar et al., 2013) and Lapointe Prostate (Lapointe et al., 2004).

#### Differential expression analysis

Differential gene and protein expression analysis was conducted using limma v.3.38.3 (Ritchie et al., 2015) R package. Limma uses linear models and borrows information across genes using empirical Bayes method and is therefore applicable for analyses of high-dimensional omics data with limited sample size. RNA-Seq data was transformed using voom. Proteomic differential expression was calculated using the algorithm for single-channel microarray gene expression data. Groups for comparison were defined in a design matrix. Linear models were fitted for expressions of each gene / intensities of each protein. Empirical Bayes method was used to borrow information across genes/proteins. Multiple testing correction was performed using the Benjamini-Hochberg method. Differential expression was defined as minimum log-FC ≥ 1 and adj. p-value ≤ 0.05.

#### Gene Set testing

For gene set testing of transcriptomic and proteomic data, the Ensemble Of Gene Set Enrichment Analyses (EGSEA) R package v.1.10.1 (Alhamdoosh et al., 2017), was used. EGSEA allows to use results from up to twelve Gene Set Enrichment (GSE)-algorithms, covering competitive and self-contained methods (Goeman and Bühlmann, 2007), to calculate collective gene set scores. We used the collective gene set score results from 11 of those methods, namely from ora, gage, camera and gsva (competitive null hypothesis) along with roast, safe, padog, plage, zscore, ssgsea and globaltest (self-contained hypothesis). We tested for enrichment on all eight Molecular Signatures Database (MSigDB) provided collections, including Gene Ontologies (GO), Kyoto Encyclopedia of Genes and Genomes (KEGG) pathways and hallmark gene sets (Liberzon et al., 2015, Liberzon et al., 2011, The Gene Ontology Consortium, 2018, Ashburner et al., 2000, Kanehisa et al., 2019, Kanehisa and Goto, 2000, Kanehisa et al., 2017, Subramanian et al., 2005). Significance was defined by an adj. p-value ≤ 0.05, adjustment method was Benjamini-Hochberg.

For overrepresentation analyses of WGCNA gene clusters and genes correlated to *STAT3* expression, please refer to the section “Weighted gene co-expression network analysis (WGCNA)” on biological theme comparison and GO enrichment of top 50 genes in the method details section.

#### Statistical analysis of targeted metabolomics data

To evaluate differences between 3 groups for 6 metabolites, ANOVA and Tukey Honest Significant Differences (HSD) test were performed for each metabolite after normality testing. Normality was tested using Pearson chi-square normality test and Levene’s test for homogeneity of variance (center = median). Visual inspection of the distribution of data was conducted using Q-Q plots and density plots. Significance was defined as p-value ≤ 0.05 after ANOVA and as adj. p-value ≤ 0.05 after TukeyHSD (95% family-wise confidence level). Since this was an exploratory experiment used for hypothesis generation, no p-value adjustment was performed between the six individual ANOVAs. Results of all ANOVAs can be found in Table S5. Statistical tests were performed using the R software environment with packages DescTools (Signorell and al., 2018) and nortest v.1.0-4 (Gross and Ligges, 2015). Plots were generated with ggpubr v.0.2 (Kassambara, 2018) and ggplot2 v.3.1.1 (Wickham, 2016.).

### DATA AND SOFTWARE AVAILABILITY

The mass spectrometry proteomics data have been deposited to the ProteomeXchange Consortium via the PRIDE (Perez-Riverol et al., 2019) partner repository with the dataset identifier PXD014251. Following publicly available datasets were used: The Cancer Genome Atlas - Prostate Adenocarcinoma (The Cancer Genome Atlas Research Network, 2015) (https://portal.gdc.cancer.gov/projects/TCGA-PRAD), Integrative genomic profiling of human prostate cancer (Taylor et al., 2010) (GSE21032), Molecular Sampling of Prostate Cancer: a dilemma for predicting disease progression (Sboner et al., 2010) (GSE16560), Gene-expression profiling of prostate tumors (Gulzar et al., 2013) (GSE40272), Gene expression profiling identifies clinically relevant subtypes of prostate cancer (Lapointe et al., 2004) (http://microarray-pubs.stanford.edu/prostateCA).

## Supplemental information

**Table S1, related to Figure 1, Figure S1 and S2. TCGA PRAD differentially expressed genes and enriched gene sets.**

**Table S2, related to Figure 2 and 3. WGCNA gene clusters.**

**Table S3, related to Figure 4. Differentially expressed proteins and enriched gene sets in human low STAT3 vs. high STAT3 FFPE sample groups.**

**Table S4, related to Figure 4. Differentially expressed proteins and enriched gene sets in *PtenStat3^pc-/-^* vs. *Pten^pc-/-^* FFPE sample groups.**

**Table S5.** related to Figure 5.

|  | ANOVA |  |  | WT-PtKO | PtSt3KO-PtKO | WT-PtSt3KO |
| --- | --- | --- | --- | --- | --- | --- |
| Metabolite | p-value | F value | df | p- adj. | p- adj. | p- adj. |
| alpha Ketoglutarate | 0.0606 | 3.761 | 2 | 0.2325 | 0.5149 | 0.0572 |
| Citrate | 0.4210 | 0.944 | 2 | 0.9737 | 0.4177 | 0.5202 |
| Fumarate | 0.0333* | 4.872 | 2 | 0.3683 | 0.1913 | 0.0268* |
| Malate | 0.0347* | 4.791 | 2 | 0.2538 | 0.2901 | 0.0292* |
| Pyruvate | 0.0097* | 7.646 | 2 | 0.0555 | 0.3728 | 0.0100* |
| Succinate | 0.0174* | 6.24 | 2 | 0.0233* | 0.9987 | 0.0512 |
| Group comparisons with Tukey multiple comparisons of means |  |  |  |  |  |  |
| 95% family-wise confidence level |  |  |  |  |  |  |

df = Degree of Freedom, WT = wild type, PtKO = *Pten<sup>pc/-</sup>*, PtSt3KO = *PtenStat3<sup>pc/-</sup>*; asterisk indicates significant p-values (< 0.05)

---

**Figure S1.**
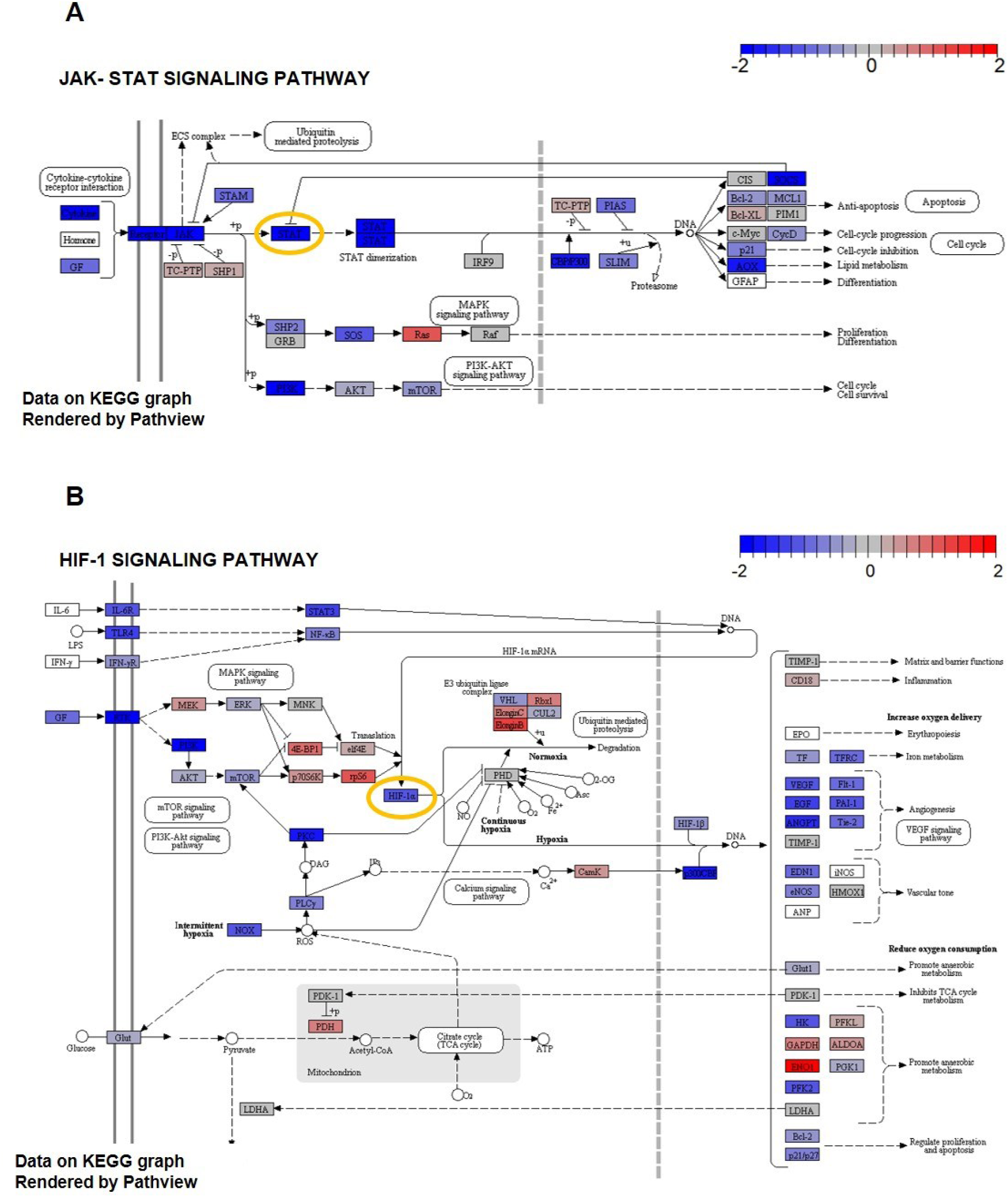
related to Figure 1 and Table S1. Scheme of downregulated KEGG pathways in low STAT3 versus high STAT3 samples. A. KEGG-representation of deregulation of the JAK-STAT signaling pathway in low STAT3 versus high STAT3 samples after testing for signaling pathways. *STAT3* is encircled in yellow. Color bar indicates z-scored deregulation of genes. Blue = downregulation, red = upregulation. B. KEGG-representation of deregulation of the HIF-1 signaling pathway in low STAT3 versus high STAT3 samples after testing for signaling pathways. *HIF-1α* is encircled in yellow. Color bar indicates z-scored deregulation of genes. Blue = downregulation, red = upregulation.

**Figure S2.**
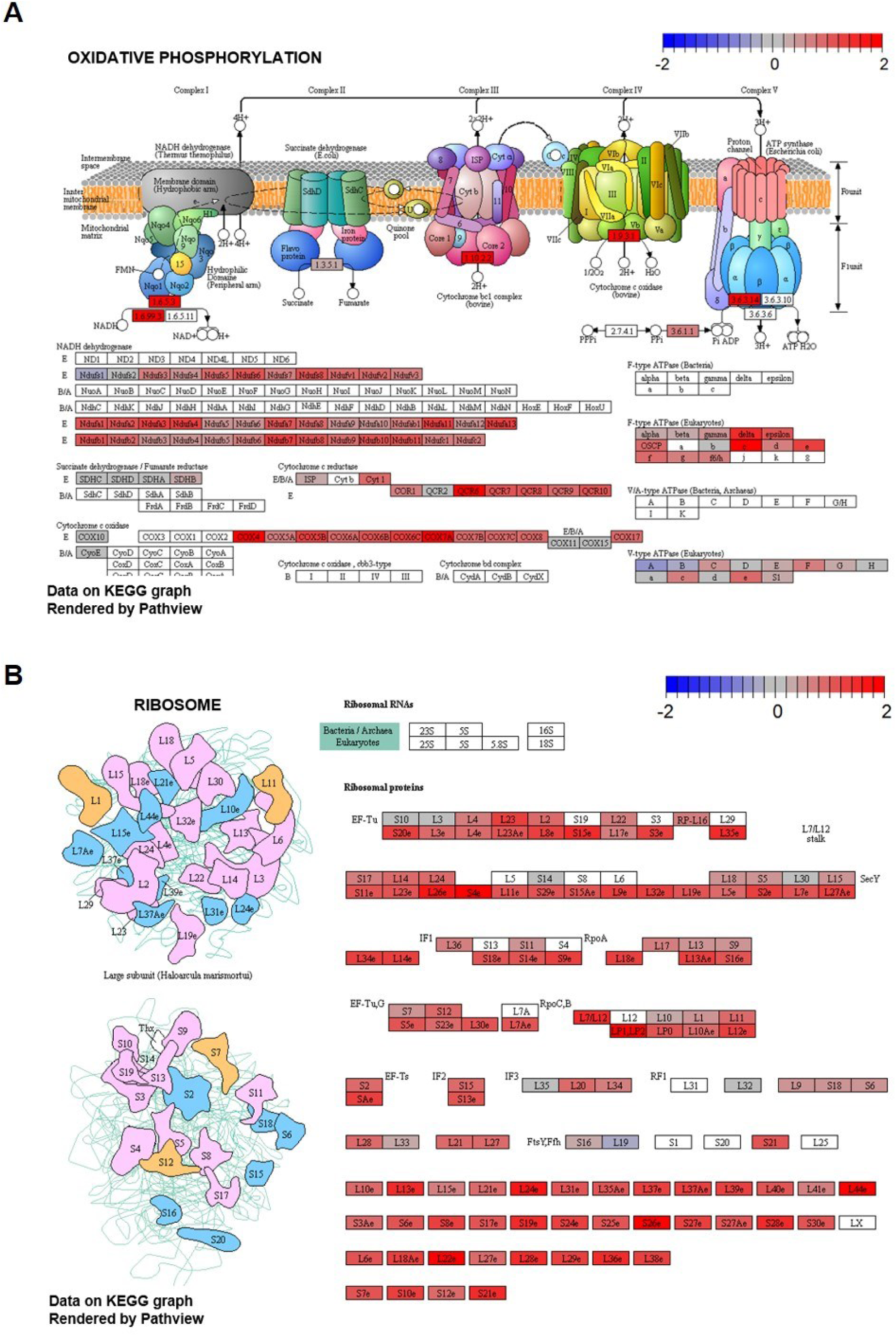
related to Figure 1 and Table S1. Scheme of upregulated KEGG pathways in low STAT3 versus high STAT3 samples. A. KEGG-representation of deregulation of oxidative phosphorylation in low STAT3 versus high STAT3 samples after testing for metabolic pathways. Color bar indicates z-scored deregulation of genes. Blue = downregulation, red = upregulation. B. KEGG-representation of deregulation of the ribosome in low STAT3 versus high STAT3 samples after testing for signaling pathways. Color bar indicates z-scored deregulation of genes. Blue = downregulation, red = upregulation.

**Figure S3.**
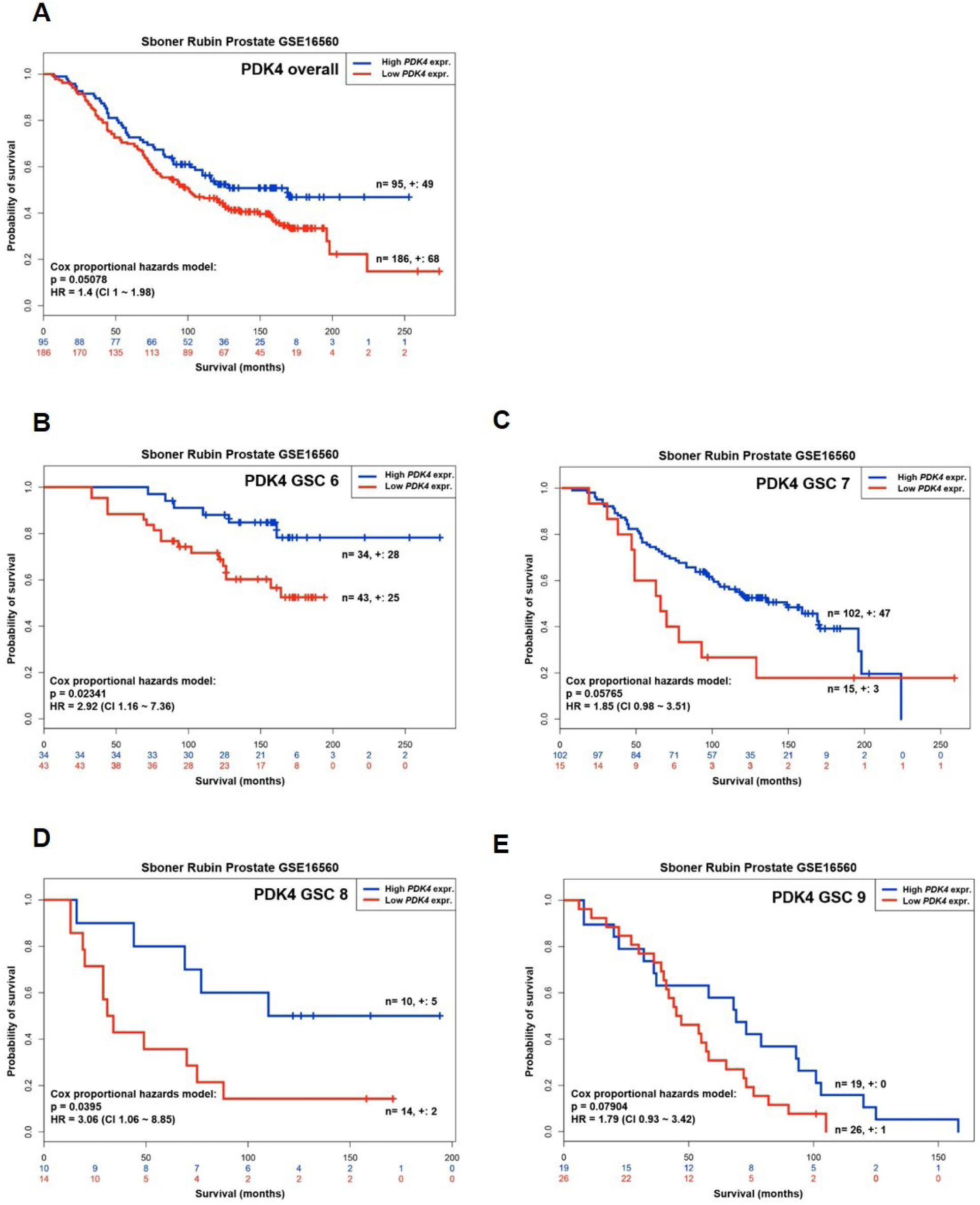
related to Figure 6. Low *PDK4* is associated with earlier death in primary PCa. A. Kaplan-Meier plot showing survival time in months for *PDK4* expression groups over all GSCs. Groups are generated by a maximizing split of samples after ranking by their prognostic index (risk score). Hazard ratio (HR), confidence intervals (CI) and p-value estimated by a Cox-model using groups as covariate are shown. Red = Low *PDK4* expression, blue = High *PDK4* expression, + = censored. B. , C. , D. and E. Kaplan-Meier plot showing survival time in months for *PDK4* expression groups for GSC 6 (B), 7 (C), 8 (D) and 9 (E). Groups are generated by a maximizing split of samples after ranking by their prognostic index (risk score). Hazard ratio (HR), confidence intervals (CI) and p-value estimated by a Cox-model using groups as covariate are shown. GSC = Gleason Score. Red = Low *PDK4* expression, blue = High *PDK4* expression, + = censored.

**Figure S4.**
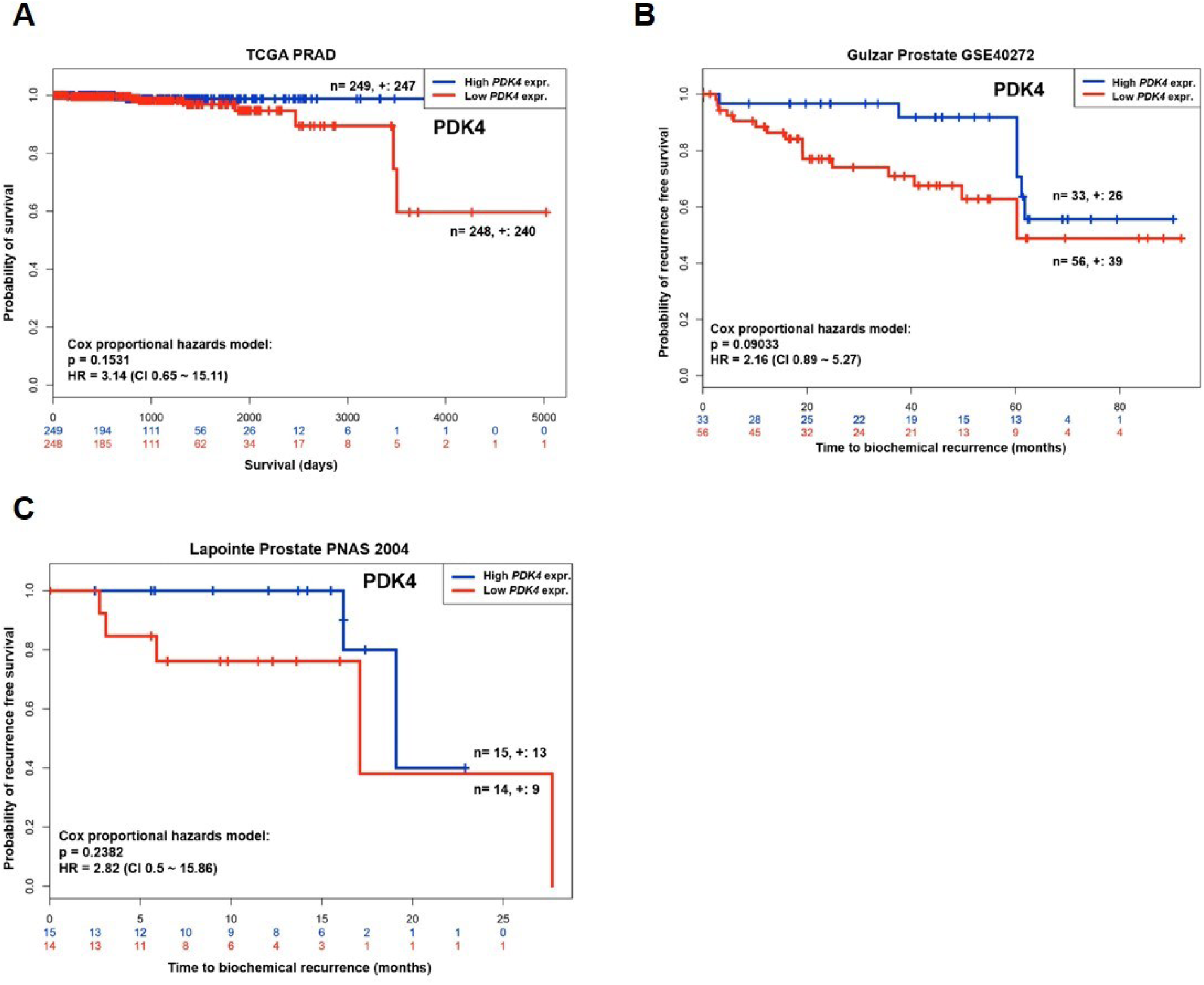
related to Figure 6. Low *PDK4* is associated with earlier biochemical recurrence or death in primary PCa. A. Kaplan-Meier plot showing survival time in days for *PDK4* expression groups in the TCGA PRAD dataset. Groups are generated by a median split of samples after ranking by their prognostic index (risk score). Hazard ratio (HR), confidence intervals (CI) and p-value estimated by a Cox-model using groups as covariate are shown. Red = Low *PDK4* expression, blue = High *PDK4* expression, + = censored. B. Kaplan-Meier plots showing time to BCR in months for *PDK4* expression groups in the Gulzar Prostate GSE40272 dataset. Groups are generated by a maximizing split of samples after ranking by their prognostic index (risk score). Hazard ratio (HR), confidence intervals (CI) and p-value estimated by a Cox-model using groups as covariate are shown. Red = Low *PDK4* expression, blue = High *PDK4* expression, + = censored. C. Kaplan-Meier plots showing time to BCR in months for *PDK4* expression groups in the Lapointe Prostate PNAS 2004 dataset. Groups are generated by a median split of samples after ranking by their prognostic index (risk score). Hazard ratio (HR), confidence intervals (CI) and p-value estimated by a Cox-model using groups as covariate are shown. Red = Low *PDK4* expression, blue = High *PDK4* expression, + = censored.

**Figure S5.**
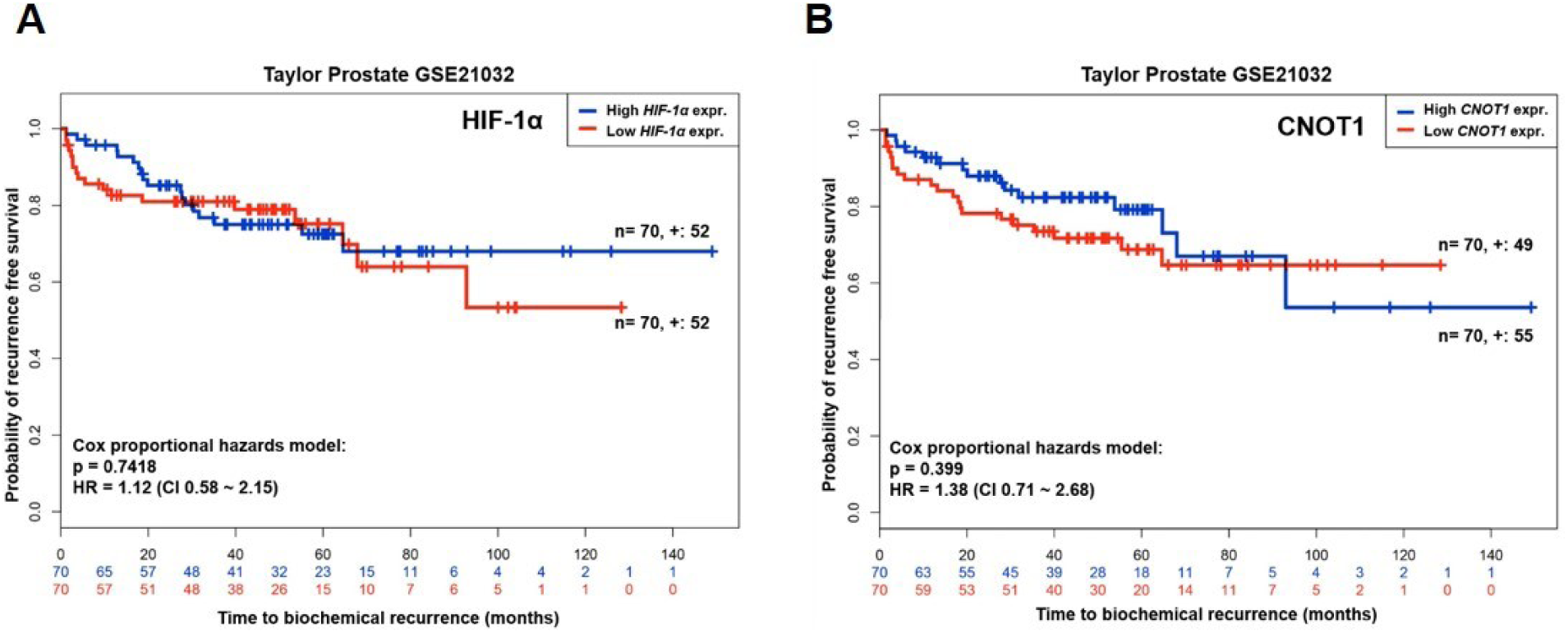
related to Figure 6. *HIF-1α* and *CNOT1* show no significant difference in time to BCR among groups. A. and B. Kaplan-Meier plots showing time to BCR in months for *HIF-1α* (A) and *CNOT1* (B) expression groups in the Taylor GSE21032 dataset. Groups are generated by a median split of samples after ranking by their prognostic index (risk score). Hazard ratio (HR), confidence intervals (CI) and p-value estimated by a Cox-model using groups as covariate are shown. Red = Low expression, blue = High expression. BCR = Biochemical recurrence.

## Exclusion of p14^ARF^/*CDKN2A* from analysis

Based on previous findings (Pencik et al., 2015) which indicate the regulation of p14^ARF^ by STAT3, resulting in more aggressive tumors after loss of both, p14^ARF^ expression was initially included in the sample groups to allow for concurrent analysis of both STAT3 and p14^ARF^.

From the TMA and patient cohort described in the Methods section, both STAT3 and p14^ARF^ protein expression were quantified by a pathologist after IHC staining (Pencik et al., 2015). Patients in the low and high STAT3 group have also low and high p14^ARF^, respectively.

Since WGCNA cluster-trait analysis for both *STAT3* and *CDKN2A* (the gene encoding for p14^ARF^) showed no influence of *CDKN2A* on respective *STAT3* correlated clusters, p14^ARF^ was neglected from further analyses and the focus set solely on STAT3.

In detail, no cluster was strongly correlated to *CDKN2A* (ρ ≤ -0.6 or ≥ 0.6, adj.p-value ≤ 0.01) after WGCNA cluster-trait analysis. Gene clusters that were moderately correlated to *CDKN2A* (ρ = ±0.3 - ±0.59, p-value ≤ 0.01) were either different to *STAT3*-correlated clusters or showed the opposite direction of correlation (data not shown). Also, *STAT3*- and *CDKN2A*-expression were not correlated with each other (ρ = -0.14, adj.p-value = 0.007, Table S2).

## References

Aguirre-Gamboa, R., Gomez-Rueda, H., Martínez-Ledesma, E., Martínez-Torteya, A., Chacolla-Huaringa, R., Rodriguez-Barrientos, A., Tamez-Peña, J. G. & Treviño, V. 2013. SurvExpress: An Online Biomarker Validation Tool and Database for Cancer Gene Expression Data Using Survival Analysis. PLOS One, 8, e74250.

Alhamdoosh, M., Ng, M., Wilson, N. J., Sheridan, J. M., Huynh, H., Wilson, M. J. & Ritchie, M. E. 2017. Combining multiple tools outperforms individual methods in gene set enrichment analyses. Bioinformatics, 33, 414–424.

Alonzi, T., Maritano, D., Gorgoni, B., Rizzuto, G., Libert, C. & Poli, V. 2001. Essential role of STAT3 in the control of the acute-phase response as revealed by inducible gene inactivation [correction of activation] in the liver. Molecular and cellular biology, 21, 1621–1632.

Anderson, N. M., Mucka, P., Kern, J. G. & Feng, H. 2018. The emerging role and targetability of the TCA cycle in cancer metabolism. Protein & cell, 9, 216–237.

Ashburner, M., Ball, C. A., Blake, J. A., Botstein, D., Butler, H., Cherry, J. M., Davis, A. P., Dolinski, K., Dwight, S. S., Eppig, J. T., Harris, M. A., Hill, D. P., Issel-Tarver, L., Kasarskis, A., Lewis, S., Matese, J. C., Richardson, J. E., Ringwald, M., Rubin, G. M. & Sherlock, G. 2000. Gene ontology: tool for the unification of biology. The Gene Ontology Consortium. Nature genetics, 25, 25–29.

Avalle, L., Camporeale, A., Morciano, G., Caroccia, N., Ghetti, E., Orecchia, V., Viavattene, D., Giorgi, C., Pinton, P. & Poli, V. 2018. STAT3 localizes to the Er, acting as a gatekeeper for Er-mitochondrion Ca2+ fluxes and apoptotic responses. Cell Death & Differentiation.

Baradaran, N., Ahmadi, H., Salem, S., Lotfi, M., Jahani, Y., Baradaran, N., Mehrsai, A. R. & Pourmand, G. 2009. The protective effect of diabetes mellitus against prostate cancer: Role of sex hormones. The Prostate, 69, 1744–1750.

Bindea, G., Mlecnik, B., Hackl, H., Charoentong, P., Tosolini, M., Kirilovsky, A., Fridman, W.-H., Pagès, F., Trajanoski, Z. & Galon, J. 2009. ClueGO: a Cytoscape plug-in to decipher functionally grouped gene ontology and pathway annotation networks. Bioinformatics (Oxford, England), 25, 1091–1093.

Birbach, A., Eisenbarth, D., Kozakowski, N., Ladenhauf, E., Schmidt-Supprian, M. & Schmid, J. A. 2011. Persistent inflammation leads to proliferative neoplasia and loss of smooth muscle cells in a prostate tumor model. Neoplasia (New York, N.Y.), 13, 692–703.

Blighe, K. 2019. EnhancedVolcano: Publication-ready volcano plots with enhanced colouring and labeling. R package version 1.0.1 ed.

Bray, F., Ferlay, J., Soerjomataram, I., Siegel, R. L., Torre, L. A. & Jemal, A. 2018. Global cancer statistics 2018: GLOBOCAN estimates of incidence and mortality worldwide for 36 cancers in 185 countries. CA: A Cancer Journal for Clinicians, 68, 394–424.

Camporeale, A., Demaria, M., Monteleone, E., Giorgi, C., Wieckowski, M. R., Pinton, P. & Poli, V. 2014. STAT3 Activities and Energy Metabolism: Dangerous Liaisons. Cancers, 6, 1579–1596.

Chen, E. Y., Tan, C. M., Kou, Y., Duan, Q., Wang, Z., Meirelles, G. V., Clark, N. R. & Ma’ayan, A. 2013. Enrichr: interactive and collaborative HTML5 gene list enrichment analysis tool. BMC Bioinformatics, 14, 128.

Chen, J., Guccini, I., di Mitri, D., Brina, D., Revandkar, A., Sarti, M., Pasquini, E., Alajati, A., Pinton, S., Losa, M., Civenni, G., Catapano, C. V., Sgrignani, J., Cavalli, A., D’antuono, R., Asara, J. M., Morandi, A., Chiarugi, P., Crotti, S., Agostini, M., Montopoli, M., Masgras, I., Rasola, A., Garcia-Escudero, R., Delaleu, N., Rinaldi, A., Bertoni, F., Bono, J. D., Carracedo, A. & Alimonti, A. 2018. Compartmentalized activities of the pyruvate dehydrogenase complex sustain lipogenesis in prostate cancer. Nature Genetics, 50, 219–228.

Choi, J. B., Moon, H. W., Park, Y. H., Bae, W. J., Cho, H. J., Hong, S.-H., Lee, J. Y., Kim, S. W., Han, K.-D. & Ha, U. S. 2016. The Impact of Diabetes on the Risk of Prostate Cancer Development according to Body Mass Index: A 10-year Nationwide Cohort Study. Journal of Cancer, 7, 2061–2066.

Colaprico, A., Silva, T. C., Olsen, C., Garofano, L., Cava, C., Garolini, D., Sabedot, T. S., Malta, T. M., Pagnotta, S. M., Castiglioni, I., Ceccarelli, M., Bontempi, G. & Noushmehr, H. 2016. TCGAbiolinks: an R/Bioconductor package for integrative analysis of TCGA data. Nucleic Acids Research, 44, e71–e71.

Costello, L. C. & Franklin, R. B. 2006. The clinical relevance of the metabolism of prostate cancer; zinc and tumor suppression: connecting the dots. Molecular cancer, 5, 17–17.

Costello, L. C., Liu, Y., Franklin, R. B. & Kennedy, M. C. 1997. Zinc Inhibition of Mitochondrial Aconitase and Its Importance in Citrate Metabolism of Prostate Epithelial Cells. Journal of Biological Chemistry, 272, 28875–28881.

Courtnay, R., Ngo, D. C., Malik, N., Ververis, K., Tortorella, S. M. & Karagiannis, T. C. 2015. Cancer metabolism and the Warburg effect: the role of Hif-1 and PI3K. Molecular Biology Reports, 42, 841–851.

Cox, J., Hein, M. Y., Luber, C. A., Paron, I., Nagaraj, N. & Mann, M. 2014. Accurate proteome-wide label-free quantification by delayed normalization and maximal peptide ratio extraction, termed MaxLFQ. Molecular & cellular proteomics: Mcp, 13, 2513–2526.

Cox, J. & Mann, M. 2008. MaxQuant enables high peptide identification rates, individualized p.p.b.-range mass accuracies and proteome-wide protein quantification. Nature Biotechnology, 26, 1367.

Cox, J., Michalski, A. & Mann, M. 2011a. Software Lock Mass by Two-Dimensional Minimization of Peptide Mass Errors. Journal of The American Society for Mass Spectrometry, 22, 1373–1380.

Cox, J., Neuhauser, N., Michalski, A., Scheltema, R. A., Olsen, J. V. & Mann, M. 2011b. Andromeda: A Peptide Search Engine Integrated into the MaxQuant Environment. Journal of Proteome Research, 10, 1794–1805.

Cutruzzolà, F., Giardina, G., Marani, M., Macone, A., Paiardini, A., Rinaldo, S. & Paone, A. 2017. Glucose Metabolism in the Progression of Prostate Cancer. Frontiers in Physiology, 8, 97.

Davis, C. A., Hitz, B. C., Sloan, C. A., Chan, E. T., Davidson, J. M., Gabdank, I., Hilton, J. A., Jain, K., Baymuradov, U. K., Narayanan, A. K., Onate, K. C., Graham, K., Miyasato, S. R., Dreszer, T. R., Strattan, J. S., Jolanki, O., Tanaka, F. Y. & Cherry, J. M. 2018. The Encyclopedia of DNA elements (ENCODE): data portal update. Nucleic Acids Res, 46, D794–d801.

Demaria, M., Giorgi, C., Lebiedzinska, M., Esposito, G., D’angeli, L., Bartoli, A., Gough, D. J., Turkson, J., Levy, D. E., Watson, C. J., Wieckowski, M. R., Provero, P., Pinton, P. & Poli, V. 2010. A STAT3-mediated metabolic switch is involved in tumour transformation and STAT3 addiction. Aging, 2, 823–842.

Donati, G., Montanaro, L. & Derenzini, M. 2012. Ribosome Biogenesis and Control of Cell Proliferation: p53 Is Not Alone. Cancer Research, 72, 1602.

Epstein, J. I., Allsbrook, W. C., Jr., Amin, M. B. & Egevad, L. L. 2005. The 2005 International Society of Urological Pathology (ISUP) Consensus Conference on Gleason Grading of Prostatic Carcinoma. Am J Surg Pathol, 29, 1228–42.

Epstein, J. I. & Lotan, T. L. 2014. The Lower Urinary Tract and Male Genital System. In: Kumar, V., Abbas, A. K. & Aster, J. C. (eds.) Robbins & Cotran Pathologic Basis of Disease. Elsevier.

Gleason, D. F. & Mellinger, G. T. 1974. Prediction of prognosis for prostatic adenocarcinoma by combined histological grading and clinical staging. J Urol, 111, 58–64.

Goeman, J. J. & Bühlmann, P. 2007. Analyzing gene expression data in terms of gene sets: methodological issues. Bioinformatics, 23, 980–987.

Gregory R. Warnes, Ben Bolker, Lodewijk Bonebakker, Robert Gentleman, Wolfgang Huber, Andy Liaw, Thomas Lumley, Martin Maechler, Arni Magnusson, Steffen Moeller, Marc Schwartz & Venables, B. 2019. gplots: Various R Programming Tools for Plotting Data. *In:* 3.0.1.1., R. P. V. (ed.). https://CRAN.R-project.org/package=gplots.

Gross, J. & Ligges, U. 2015. nortest: Tests for Normality.: R package version 1.0-4.

Guda, M. R., Asuthkar, S., Labak, C. M., Tsung, A. J., Alexandrov, I., Mackenzie, M. J., Prasad, D. V. & Velpula, K. K. 2018. Targeting PDK4 inhibits breast cancer metabolism. American journal of cancer research, 8, 1725–1738.

Gulzar, Z. G., Mckenney, J. K. & Brooks, J. D. 2013. Increased expression of NuSAP in recurrent prostate cancer is mediated by E2F1. Oncogene, 32, 70–77.

Hanahan, D. & Weinberg, Robert A. 2011. Hallmarks of Cancer: The Next Generation. Cell, 144, 646–674.

Huynh, J., Chand, A., Gough, D. & Ernst, M. 2019. Therapeutically exploiting STAT3 activity in cancer — using tissue repair as a road map. Nature Reviews Cancer, 19, 82–96.

Iglesias-Gato, D., Wikström, P., Tyanova, S., Lavallee, C., Thysell, E., Carlsson, J., Hägglöf, C., Cox, J., Andrén, O., Stattin, P., Egevad, L., Widmark, A., Bjartell, A., Collins, C. C., Bergh, A., Geiger, T., Mann, M. & Flores-Morales, A. 2016. The Proteome of Primary Prostate Cancer. European Urology, 69, 942–952.

Jeoung, N. H. 2015. Pyruvate Dehydrogenase Kinases: Therapeutic Targets for Diabetes and Cancers. Diabetes & metabolism journal, 39, 188–197.

Kanehisa, M., Furumichi, M., Tanabe, M., Sato, Y. & Morishima, K. 2017. KEGG: new perspectives on genomes, pathways, diseases and drugs. Nucleic Acids Res, 45, D353–d361.

Kanehisa, M. & Goto, S. 2000. KEGG: kyoto encyclopedia of genes and genomes. Nucleic acids research, 28, 27–30.

Kanehisa, M., Sato, Y., Furumichi, M., Morishima, K. & Tanabe, M. 2019. New approach for understanding genome variations in KEGG. Nucleic Acids Res, 47, D590–d595.

Kassambara, A. 2018. ggpubr: ‘ggplot2’ Based Publication Ready Plots.: R package version 0.2.

Kassambara, A., Kosinski, M. & Biecek, P. 2019. survminer: Drawing Survival Curves using ‘ggplot2’. R package version 0.4.5. Available from: https://CRAN.R-project.org/package=survminer.

Kim, Y., Jeon, J., Mejia, S., Yao, C. Q., Ignatchenko, V., Nyalwidhe, J. O., Gramolini, A. O., Lance, R. S., Troyer, D. A., Drake, R. R., Boutros, P. C., Semmes, O. J. & Kislinger, T. 2016. Targeted proteomics identifies liquid-biopsy signatures for extracapsular prostate cancer. Nature Communications, 7, 11906.

Kuleshov, M. V., Jones, M. R., Rouillard, A. D., Fernandez, N. F., Duan, Q., Wang, Z., Koplev, S., Jenkins, S. L., Jagodnik, K. M., Lachmann, A., Mcdermott, M. G., Monteiro, C. D., Gundersen, G. W. & Ma’ayan, A. 2016. Enrichr: a comprehensive gene set enrichment analysis web server 2016 update. Nucleic Acids Res, 44, W90–7.

Kulkarni, S. S., Salehzadeh, F., Fritz, T., Zierath, J. R., Krook, A. & Osler, M. E. 2012. Mitochondrial regulators of fatty acid metabolism reflect metabolic dysfunction in type 2 diabetes mellitus. Metabolism, 61, 175–185.

Langfelder, P. & Horvath, S. 2008. WGCNA: an R package for weighted correlation network analysis. BMC Bioinformatics, 9, 559.

Langfelder, P. & Horvath, S. 2012. Fast R Functions for Robust Correlations and Hierarchical Clustering. Journal of Statistical Software; Vol 1, Issue 11 (2012).

Lapointe, J., Li, C., Higgins, J. P., van de Rijn, M., Bair, E., Montgomery, K., Ferrari, M., Egevad, L., Rayford, W., Bergerheim, U., Ekman, P., Demarzo, A. M., Tibshirani, R., Botstein, D., Brown, P. O., Brooks, J. D. & Pollack, J. R. 2004. Gene expression profiling identifies clinically relevant subtypes of prostate cancer. Proceedings of the National Academy of Sciences of the United States of America, 101, 811.

Latonen, L., Afyounian, E., Jylhä, A., Nättinen, J., Aapola, U., Annala, M., Kivinummi, K. K., Tammela, T. T. L., Beuerman, R. W., Uusitalo, H., Nykter, M. & Visakorpi, T. 2018. Integrative proteomics in prostate cancer uncovers robustness against genomic and transcriptomic aberrations during disease progression. Nature Communications, 9, 1176.

Law, C. W., Chen, Y., Shi, W. & Smyth, G. K. 2014. voom: precision weights unlock linear model analysis tools for Rna-seq read counts. Genome Biology, 15, R29.

Lee, I.-K. 2014. The role of pyruvate dehydrogenase kinase in diabetes and obesity. Diabetes & metabolism journal, 38, 181–186.

Liberzon, A., Birger, C., Thorvaldsdóttir, H., Ghandi, M., Mesirov, Jill P. & Tamayo, P. 2015. The Molecular Signatures Database Hallmark Gene Set Collection. Cell Systems, 1, 417–425.

Liberzon, A., Subramanian, A., Pinchback, R., Thorvaldsdóttir, H., Tamayo, P. & Mesirov, J. P. 2011. Molecular signatures database (MSigDB) 3.0. Bioinformatics, 27, 1739–1740.

Liu, Z., Chen, X., Wang, Y., Peng, H., Wang, Y., Jing, Y. & Zhang, H. 2014. PDK4 protein promotes tumorigenesis through activation of camp-response element-binding protein (CREB)-Ras homolog enriched in brain (RHEB)-mTORC1 signaling cascade. J Biol Chem, 289, 29739–49.

Mengual, L., Ars, E., Lozano, J. J., Burset, M., Izquierdo, L., Ingelmo-Torres, M., Gaya, J. M., Algaba, F., Villavicencio, H., Ribal, M. J. & Alcaraz, A. 2014. Gene expression profiles in prostate cancer: identification of candidate non-invasive diagnostic markers. Actas Urol Esp, 38, 143–9.

National Collaborating Centre for Cancer 2014. Prostate Cancer: Diagnosis and Treatment. NICE Clinical Guidelines, No. 175. Cardiff (UK): National Collaborating Centre for Cancer (UK);.

Niu, G., Briggs, J., Deng, J., Ma, Y., Lee, H., Kortylewski, M., Kujawski, M., Kay, H., Cress, W. D., Jove, R. & Yu, H. 2008. Signal transducer and activator of transcription 3 is required for hypoxia-inducible factor-1alpha RNA expression in both tumor cells and tumor-associated myeloid cells. Molecular cancer research: Mcr, 6, 1099–1105.

Ostasiewicz, P., Zielinska, D. F., Mann, M. & Wisniewski, J. R. 2010. Proteome, phosphoproteome, and N-glycoproteome are quantitatively preserved in formalin-fixed paraffin-embedded tissue and analyzable by high-resolution mass spectrometry. J Proteome Res, 9, 3688–700.

Pawlus, M. R., Wang, L. & Hu, C. J. 2014. STAT3 and HIF1α cooperatively activate HIF1 target genes in Mda-Mb-231 and RCC4 cells. Oncogene, 33, 1670–1679.

Pencik, J., Schlederer, M., Gruber, W., Unger, C., Walker, S. M., Chalaris, A., Marié, I. J., Hassler, M. R., Javaheri, T., Aksoy, O., Blayney, J. K., Prutsch, N., Skucha, A., Herac, M., Krämer, O. H., Mazal, P., Grebien, F., Egger, G., Poli, V., Mikulits, W., Eferl, R., Esterbauer, H., Kennedy, R., Fend, F., Scharpf, M., Braun, M., Perner, S., Levy, D. E., Malcolm, T., Turner, S. D., Haitel, A., Susani, M., Moazzami, A., Rose-John, S., Aberger, F., Merkel, O., Moriggl, R., Culig, Z., Dolznig, H. & Kenner, L. 2015. STAT3 regulated ARF expression suppresses prostate cancer metastasis. Nature Communications, 6, 7736.

Perez-Riverol, Y., Csordas, A., Bai, J., Bernal-Llinares, M., Hewapathirana, S., Kundu, D. J., Inuganti, A., Griss, J., Mayer, G., Eisenacher, M., Perez, E., Uszkoreit, J., Pfeuffer, J., Sachsenberg, T., Yilmaz, S., Tiwary, S., Cox, J., Audain, E., Walzer, M., Jarnuczak, A. F., Ternent, T., Brazma, A. & Vizcaino, J. A. 2019. The PRIDE database and related tools and resources in 2019: improving support for quantification data. Nucleic Acids Res, 47, D442–d450.

Pohlert, T. 2018. PMCMRplus: Calculate Pairwise Multiple Comparisons of Mean Rank Sums Extended.: R package version 1.4.1.

Poli, V. & Camporeale, A. 2015. STAT3-Mediated Metabolic Reprograming in Cellular Transformation and Implications for Drug Resistance. Frontiers in Oncology, 5, 121.

Ritchie, M. E., Phipson, B., Wu, D., Hu, Y., Law, C. W., Shi, W. & Smyth, G. K. 2015. limma powers differential expression analyses for Rna-sequencing and microarray studies. Nucleic Acids Res, 43, e47.

Robinson, M. D., Mccarthy, D. J. & Smyth, G. K. 2010. edgeR: a Bioconductor package for differential expression analysis of digital gene expression data. Bioinformatics, 26, 139–40.

Robinson, M. D. & Oshlack, A. 2010. A scaling normalization method for differential expression analysis of Rna-seq data. Genome Biology, 11, R25.

Roulhac, P. L., Ward, J. M., Thompson, J. W., Soderblom, E. J., Silva, M., Moseley, M. A., 3rd & Jarvis, E. D. 2011. Microproteomics: quantitative proteomic profiling of small numbers of laser-captured cells. Cold Spring Harbor protocols, 2011, pdb.prot5573-pdb.prot5573.

Sathianathen, N. J., Konety, B. R., Crook, J., Saad, F. & Lawrentschuk, N. 2018. Landmarks in prostate cancer. Nature Reviews Urology, 15, 627–642.

Sboner, A., Demichelis, F., Calza, S., Pawitan, Y., Setlur, S. R., Hoshida, Y., Perner, S., Adami, H.-O., Fall, K., Mucci, L. A., Kantoff, P. W., Stampfer, M., Andersson, S.-O., Varenhorst, E., Johansson, J.-E., Gerstein, M. B., Golub, T. R., Rubin, M. A. & Andrén, O. 2010. Molecular sampling of prostate cancer: a dilemma for predicting disease progression. BMC Medical Genomics, 3, 8.

Schaab, C., Geiger, T., Stoehr, G., Cox, J. & Mann, M. 2012. Analysis of high accuracy, quantitative proteomics data in the MaxQB database. Molecular & cellular proteomics: Mcp, 11, M111.014068-M111.014068.

Schwaiger, M., Schoeny, H., El Abiead, Y., Hermann, G., Rampler, E. & Koellensperger, G. 2019. Merging metabolomics and lipidomics into one analytical run. Analyst, 144, 220–229.

Signorell, A. & Al., E. M. 2018. DescTools: Tools for descriptive statistics.: R package version 0.99.26.

Sinha, A., Huang, V., Livingstone, J., Wang, J., Fox, N. S., Kurganovs, N., Ignatchenko, V., Fritsch, K., Donmez, N., Heisler, L. E., Shiah, Y.-J., Yao, C. Q., Alfaro, J. A., Volik, S., Lapuk, A., Fraser, M., Kron, K., Murison, A., Lupien, M., Sahinalp, C., Collins, C. C., Tetu, B., Masoomian, M., Berman, D. M., van der Kwast, T., Bristow, R. G., Kislinger, T. & Boutros, P. C. 2019. The Proteogenomic Landscape of Curable Prostate Cancer. Cancer Cell, 35, 414–427.e6.

Stacpoole, P. W. 2017. Therapeutic Targeting of the Pyruvate Dehydrogenase Complex/Pyruvate Dehydrogenase Kinase (PDC/PDK) Axis in Cancer. JNCI: Journal of the National Cancer Institute, 109.

Steitz, T. A. 2008. A structural understanding of the dynamic ribosome machine. Nature Reviews Molecular Cell Biology, 9, 242.

Stelzer, G., Rosen, N., Plaschkes, I., Zimmerman, S., Twik, M., Fishilevich, S., Stein, T. I., Nudel, R., Lieder, I., Mazor, Y., Kaplan, S., Dahary, D., Warshawsky, D., Guan-Golan, Y., Kohn, A., Rappaport, N., Safran, M. & Lancet, D. 2016. The GeneCards Suite: From Gene Data Mining to Disease Genome Sequence Analyses. Current Protocols in Bioinformatics, 54, 1.30.1-1.30.33.

Subramanian, A., Tamayo, P., Mootha, V. K., Mukherjee, S., Ebert, B. L., Gillette, M. A., Paulovich, A., Pomeroy, S. L., Golub, T. R., Lander, E. S. & Mesirov, J. P. 2005. Gene set enrichment analysis: A knowledge-based approach for interpreting genome-wide expression profiles. Proceedings of the National Academy of Sciences, 102, 15545–15550.

Sun, Y., Daemen, A., Hatzivassiliou, G., Arnott, D., Wilson, C., Zhuang, G., Gao, M., Liu, P., Boudreau, A., Johnson, L. & Settleman, J. 2014. Metabolic and transcriptional profiling reveals pyruvate dehydrogenase kinase 4 as a mediator of epithelial-mesenchymal transition and drug resistance in tumor cells. Cancer & Metabolism, 2, 20.

Suzuki, A., Yamaguchi, M. T., Ohteki, T., Sasaki, T., Kaisho, T., Kimura, Y., Yoshida, R., Wakeham, A., Higuchi, T., Fukumoto, M., Tsubata, T., Ohashi, P. S., Koyasu, S., Penninger, J. M., Nakano, T. & Mak, T. W. 2001. T Cell-Specific Loss of Pten Leads to Defects in Central and Peripheral Tolerance. Immunity, 14, 523–534.

Taylor, B. S., Schultz, N., Hieronymus, H., Gopalan, A., Xiao, Y., Carver, B. S., Arora, V. K., Kaushik, P., Cerami, E., Reva, B., Antipin, Y., Mitsiades, N., Landers, T., Dolgalev, I., Major, E., Wilson, M., Socci, N. D., Lash, A. E., Heguy, A., Eastham, J. A., Scher, H. I., Reuter, V. E., Scardino, P. T., Sander, C., Sawyers, C. L. & Gerald, W. L. 2010. Integrative Genomic Profiling of Human Prostate Cancer. Cancer Cell, 18, 11–22.

The Cancer Genome Atlas Research Network 2015. The Molecular Taxonomy of Primary Prostate Cancer. Cell, 163, 1011–1025.

The Gene Ontology Consortium 2018. The Gene Ontology Resource: 20 years and still GOing strong. Nucleic Acids Research, 47, D330–D338.

Therneau, T. M. 2015. A Package for Survival Analysis in S. version 2.38.

Therneau, T. M. & Grambsch, P. M. 2000. Modeling Survival Data: Extending the Cox Model, New York, Springer.

Tyanova, S., Temu, T., Carlson, A., Sinitcyn, P., Mann, M. & Cox, J. 2015. Visualization of Lc-MS/MS proteomics data in MaxQuant. Proteomics, 15, 1453–1456.

Tyanova, S., Temu, T. & Cox, J. 2016a. The MaxQuant computational platform for mass spectrometry-based shotgun proteomics. Nat. Protocols, 11, 2301–2319.

Tyanova, S., Temu, T., Sinitcyn, P., Carlson, A., Hein, M. Y., Geiger, T., Mann, M. & Cox, J. 2016b. The Perseus computational platform for comprehensive analysis of (prote)omics data. Nat Meth, 13, 731–740.

Wang, H., Qian, W.-J., Mottaz, H. M., Clauss, T. R. W., Anderson, D. J., Moore, R. J., Camp, D. G., 2nd, Khan, A. H., Sforza, D. M., Pallavicini, M., Smith, D. J. & Smith, R. D. 2005. Development and evaluation of a micro- and nanoscale proteomic sample preparation method. Journal of proteome research, 4, 2397-2403.

Wegrzyn, J., Potla, R., Chwae, Y.-J., Sepuri, N. B. V., Zhang, Q., Koeck, T., Derecka, M., Szczepanek, K, Szelag, M., Gornicka, A., Moh, A., Moghaddas, S., Chen, Q., Bobbili, S., Cichy, J., Dulak, J., Baker, D. P., Wolfman, A., Stuehr, D., Hassan, M. O., Fu, X.-Y., Avadhani, N., Drake, J. I., Fawcett, P., Lesnefsky, E. J. & Larner, A. C. 2009. Function of Mitochondrial Stat3 in Cellular Respiration. Science, 323, 793.

White, U. A., Coulter, A. A., Miles, T. K. & Stephens, J. M. 2007. The STAT5A-Mediated Induction of Pyruvate Dehydrogenase Kinase 4 Expression by Prolactin or Growth Hormone in Adipocytes. Diabetes, 56, 1623.

Wickham, H. (ed.) 2016. ggplot2: Elegant Graphics for Data Analysis., New York: Springer-Verlag.

Wickham, H. 2017. tidyverse: Easily Install and Load the ‘Tidyverse’. R package version1.2.1.

Wisniewski, J. R., Dus, K. & Mann, M. 2013. Proteomic workflow for analysis of archival formalin-fixed and paraffin-embedded clinical samples to a depth of 10 000 proteins. Proteomics Clin Appl, 7, 225–33.

Woolbright, B. L., Choudhary, D., Mikhalyuk, A., Trammel, C., Shanmugam, S., Abbott, E., Pilbeam, C. C. & Taylor, J. A. 2018. The Role of Pyruvate Dehydrogenase Kinase-4 (PDK4) in Bladder Cancer and Chemoresistance. Molecular Cancer Therapeutics, 17, 2004.

Wu, X., Wu, J., Huang, J., Powell, W. C., Zhang, J., Matusik, R. J., Sangiorgi, F. O., Maxson, R. E., Sucov, H. M. & Roy-Burman, P. 2001. Generation of a prostate epithelial cell-specific Cre transgenic mouse model for tissue-specific gene ablation. Mechanisms of Development, 101, 61–69.

Yamane, K., Indalao, I. L., Chida, J., Yamamoto, Y., Hanawa, M. & Kido, H. 2014. Diisopropylamine Dichloroacetate, a Novel Pyruvate Dehydrogenase Kinase 4 Inhibitor, as a Potential Therapeutic Agent for Metabolic Disorders and Multiorgan Failure in Severe Influenza. PLOS One, 9, e98032.

Yang, C., Wang, S., Ruan, H., Li, B., Cheng, Z., He, J., Zuo, Q., Yu, C., Wang, H., Lv, Y., Gu, D., Jin, G., Yao, M., Qin, W. & Jin, H. 2019. Downregulation of PDK4 Increases Lipogenesis and Associates with Poor Prognosis in Hepatocellular Carcinoma. J Cancer, 10, 918–926.

Yip, A. M. & Horvath, S. 2007. Gene network interconnectedness and the generalized topological overlap measure. BMC Bioinformatics, 8, 22.

Yu, G., Wang, L.-G., Han, Y. & He, Q.-Y. 2012. clusterProfiler: an R Package for Comparing Biological Themes Among Gene Clusters. OMICS: A Journal of Integrative Biology, 16, 284–287.

Zhang, B. & Horvath, S. 2005. A general framework for weighted gene co-expression network analysis. Stat Appl Genet Mol Biol, 4, Article17.

Zhang, S., Hulver, M. W., Mcmillan, R. P., Cline, M. A. & Gilbert, E. R. 2014. The pivotal role of pyruvate dehydrogenase kinases in metabolic flexibility. Nutrition & Metabolism, 11, 10.

